# A Population-Scale Single-Cell Atlas of the Human Heart Reveals Cellular Remodeling and Cell–Cell Communication in Aging and Cardiac Disease

**DOI:** 10.64898/2025.12.12.693831

**Authors:** Zheng Gong, Mengwei Li, Kok Siong Ang, Xi Zhao, Chrishan J Ramachandra, Xiaomeng Wang, Derek John Hausenloy, Yibin Wang, Yuguo Chen, Jinmiao Chen

## Abstract

Previous single-cell and single-nucleus heart atlases, often limited by small sample sizes, lack the statistical power needed for phenotype association analysis, particularly for cardiovascular diseases and cardiac aging. To address this, we integrated data from 436 samples across 12 single-cell studies, harmonized the corresponding sample metadata, and constructed a comprehensive heart atlas comprising 355,762 cells and 1,436,719 nuclei. Consensus annotation identified 10 broad cell types and 54 fine-grained subsets. Associating gene expression patterns and cell type proportions with phenotypic data, we identified *NRG1*-expressing endocardial cells linked to multiple cardiac diseases and found that interferon (IFN) response signatures mark aging in multiple heart cell types. Importantly, we also developed PopComm, a novel computational method for inferring ligand–receptor (LR) interactions from population-scale single-cell data and quantifying interaction strength for individual samples. Using PopComm, we revealed a close association between the IFN response state and altered cell–cell communication during cardiac aging.

## Main

The human heart performs central roles in circulation, oxygenation, and nutrient delivery^1,2^. It accomplishes these physiological functions through the synchronized actions of the different types of specialized cardiac cells^3^. Together, these diverse cell types ensure the heart’s proper function. Critically, these cells do not act in isolation; they communicate continuously through intricate ligand–receptor signaling and physical interactions. These communications are essential for coordinating heart development, maintaining tissue homeostasis, and responding to stress or injury^4–6^. During aging, the cardiac cells and their intercellular communications undergo quantitative and qualitative alterations. Cardiomyocytes often decrease in number and may exhibit mitochondrial dysfunction or hypertrophy^7,8^. Fibroblasts shift toward a pro-fibrotic and pro-inflammatory secretory phenotype, while endothelial cells (ECs) increasingly exhibit senescent traits, impairing vascular integrity and angiogenesis^9–11^. A central hallmark of cardiac aging is the accumulation of senescent cells that secrete a range of inflammatory cytokines, growth factors, and matrix remodeling enzymes, a phenomenon termed the senescence-associated secretory phenotype (SASP)^9^. Together with the compromised roles of non-myocytes, these changes synergistically drive the progression of cardiac aging^12^.

Cardiac aging and related cardiovascular diseases burden global health and economies, reinforcing the urgent need to clarify their cellular and molecular mechanisms^13^. For tackling this, single-cell technologies hold great promise for unraveling the intricate interplay between cell types, molecular pathways, and physiological processes in cardiac aging and diseases^14,15^. Indeed, existing scRNA-seq and snRNA-seq studies on the heart have provided valuable insights into cardiac cellular heterogeneity, gene expression profiles, and regulatory networks^3,16–38^. These studies have characterized diverse cell populations within the heart, revealing their distinct molecular signatures and functional roles in cardiac physiology and pathology (**summarized in Supplementary Table 1**).

Despite these advancements, existing studies often encounter limitations due to small sample sizes, which hinder the study of rare cell types and phenotype-related changes at the cellular level^39^. Therefore, accurate and sensitive identification of phenotype-related cell types and their gene expression change requires larger sample sizes and integrated datasets from single-cell and single-nucleus RNA-seq studies. Another major limitation is the lack of tools for analyzing cell–cell communication at the population level. Existing methods like CellPhoneDB^40^ and CellChat^41^ perform well for individual datasets or aggregated samples but do not provide communication scores at the sample level, making it difficult to associate intercellular signaling changes with phenotypic variation.

In this study, we address these limitations by integrating a large number of samples with well-curated metadata to construct a comprehensive population-scale single-cell and single-nucleus RNA-seq heart atlas. Powered by the large sample size and high-quality metadata, our phenotype association analyses identified Neuregulin-1 (*NRG1*)-expressing endocardial cells associated with multiple cardiovascular diseases, implicating them in fibrotic remodeling and pathological progression. Additionally, we observed a pronounced shift in cardiac cellular and molecular states after the age of 60, suggesting a pivotal aging breakpoint. Notably, interferon-related inflammation emerges as a hallmark of cardiac aging, with broad impacts across multiple cell types.

We also developed PopComm (a framework for population-level cell–cell communication) for inferring and quantifying ligand–receptor (LR) interactions from large-scale single-cell datasets. Unlike existing tools, PopComm quantifies the strength of specific LR interactions at the sample level, allowing direct correlation with phenotypic traits such as age, disease status, and immune response. Applying PopComm to our heart atlases, we found that heart disease markedly reshapes the LR interaction landscape, with interactions involving cardiomyocytes and endocardial cells playing important roles in disease-associated signaling. In aging, we discovered that changes in intercellular communication were more strongly correlated with interferon (IFN) response states than with chronological age. Together, these findings highlight the utility of PopComm in uncovering biologically meaningful and phenotype-associated cell–cell communication patterns. By enabling sample-level resolution and phenotype integration, PopComm provides a powerful framework for studying cell–cell interactions during aging, disease progression, and immune-mediated remodeling.

## Results

### Mapping cardiac lineages with single-cell and single-nucleus transcriptomics

Understanding the cellular composition of the human heart is essential for uncovering how it develops, adapts to stress, and undergoes disease- or aging-related decline. While individual single-cell or single-nucleus studies have revealed important aspects of cardiac biology, they are typically limited in scope due to focus on one disease, age group, or heart region^3,17–20,22,24,26,28,29,34,36^. To overcome these limitations, we sought to construct a large-scale single-cell heart atlas from public data. We first systematically assembled scRNA-seq and snRNA-seq data from 459 (before quality control) non-fetal samples in the DISCO database^42,43^. Following stringent project-level quality control, 355,762 cells and 1,436,719 nuclei from 436 samples were retained for further analysis. The comprehensive metadata, including tissue region, disease, sex, and age of these samples, were manually curated from the original studies (**Fig. 1a; Supplementary Table 2**). The collected samples encompass six different regions of the heart, namely the left atrium (LA), right atrium (RA), left ventricle (LV), right ventricle (RV), apex (AP), and septum (S). In terms of condition, the samples consist of both normal controls and eight disease types. The samples also span a wide age range, from preschoolers (0–9 years) to older adults (70–79 years), capturing developmental, aging-related and diseases associated changes. This comprehensive dataset provides the breadth needed to chart cardiac cell states across development, aging, and disease.

**Fig. 1:**
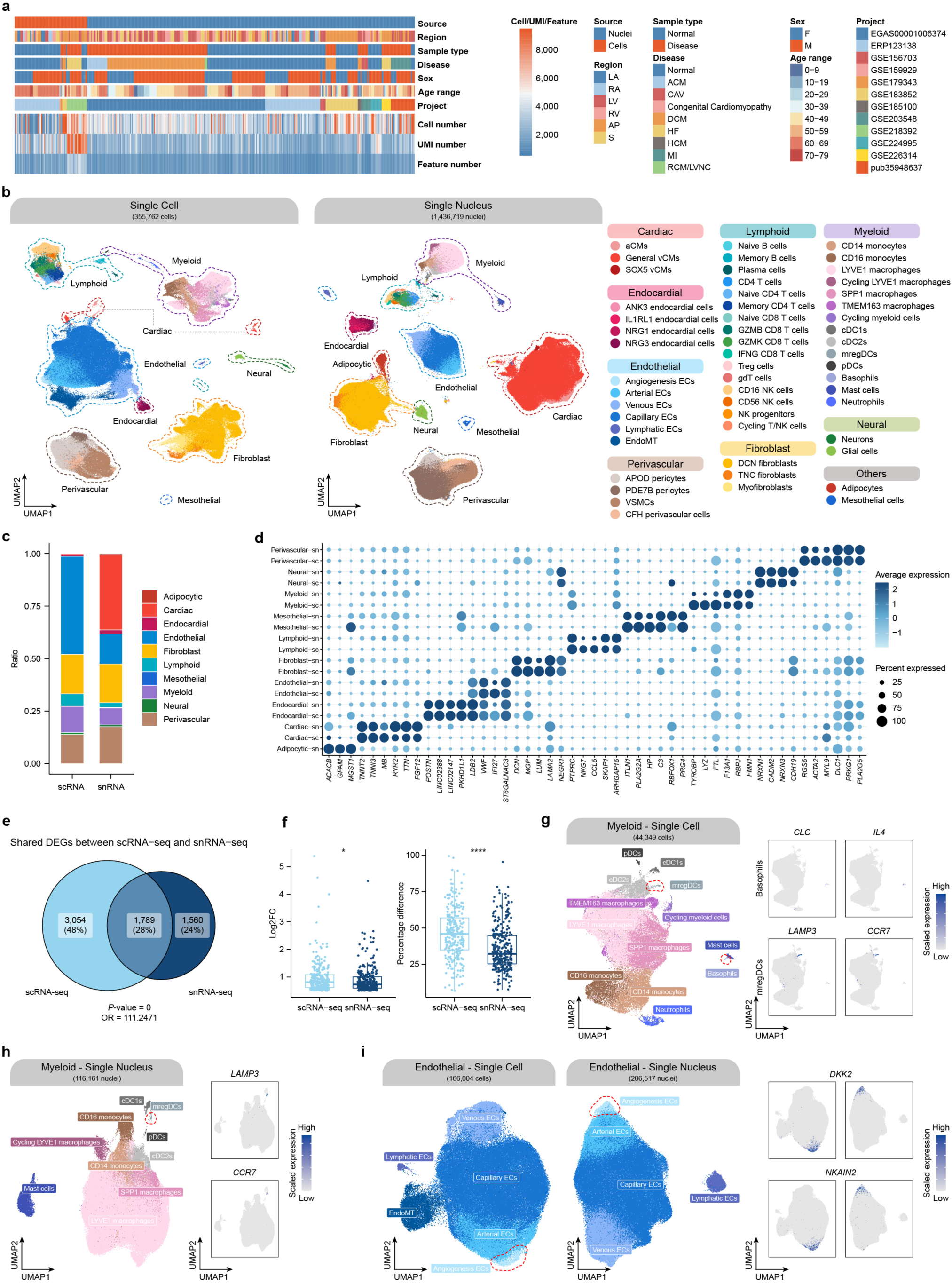
Development of comprehensive single-cell and single-nuclei atlas of heart. **a**, Overview of sample metadata and quality metrics. **b**, UMAP representations of integrated scRNA-seq and snRNA-seq data. **c**, The proportion of main cell types in scRNA-seq and snRNA-seq data. **d**, Dot plot representation of top marker genes in scRNA-seq and snRNA-seq data. Top 3 markers ordered by fold change in each cell type and each modality are shown. **e**, Venn diagram of unique and shared marker genes identified in scRNA-seq and snRNA-seq data. **f**, Box plots showing the log2 fold change (Log2FC, left) and percentage difference (right) in marker gene expression between scRNA-seq and snRNA-seq data. **g**, Basophils, mregDCs and marker gene expression in myeloid subsets from scRNA-seq data. UMAP visualization of basophils and mregDCs (left), and gene expression of *CLC*, *IL4*, *LAMP3*, and *CCR7* (right) in myeloid subsets. **h**, mregDCs and marker gene expression in the myeloid subsets from snRNA-seq data. UMAP visualization of mregDCs (left) and gene expression of *LAMP3* and *CCR7* (right) in myeloid subsets. **i**, Angiogenesis ECs and marker gene expression in endothelial subsets. UMAP visualization of Angiogenesis ECs from scRNA-seq and snRNA-seq data (left) and gene expression of *DKK2* and *NKAIN2* across endothelial subsets (right). Significance (two-sided Wilcoxon test) is indicated: \**P* < 0.05, \*\*\*\**P* < 0.0001.

As scRNA-seq and snRNA-seq capture different transcript pools and show differential sensitivities in detecting distinct cell types, we treated them as different modalities for separate data integration. Merging them directly risks conflating technical and biological variations, whereas separate integration preserves complementary signals and enables comprehensive mapping of cardiac cell states. For each modality, we applied hierarchical integration with scVI^44^ independently. The first round of integration, followed by unsupervised clustering, identified 10 major cell types: cardiomyocytes, endocardial cells, endothelial cells, perivascular cells, lymphoid cells, myeloid cells, fibroblasts, mesothelial cells, neural cells, and adipocytes (**Fig. 1b**). Notably, scRNA-seq and snRNA-seq exhibited distinct biases in the recovered cell populations (**Fig. 1c**). For instance, adipocytes were captured by snRNA-seq but not by scRNA-seq. Similarly, cardiomyocytes were the most abundant cell type in the snRNA-seq data (35.9% of cells) but comprised only 0.6% of the scRNA-seq data. Endothelial cell abundance was also notably different, constituting 14.4% of the snRNA-seq but 46.7% of scRNA-seq.

We then computed the differentially expressed genes (DEGs) in both modalities to facilitate cell type annotation. The top DEGs (**Supplementary Tables 5 & 8**) identified for each major cell type were consistent with previously reported markers (**Fig. 1d**), highlighting the reliability of our annotation. A systematic comparison of DEGs between the two modalities revealed that 27.9% of DEGs were shared, achieving an odds ratio (OR) of 111 with a statistically significant *P*-value (*P*-value = 0, OR = 111.25; **Fig. 1e)**. Additionally, comparisons of log2 fold changes (log2FC) and percentage differences showed that the DEGs of scRNA-seq and snRNA-seq exhibited similar log2FC values, but the scRNA-seq analysis produced significantly higher percentage differences (**Fig. 1f**). This finding indicates differences in detection sensitivity between the two techniques, particularly for certain cell types and their marker gene expression profiles, with the scRNA-seq profiles showing a greater dynamic range of gene expression.

To resolve fine-grained cell types, we performed a second round of integration within each major cell type (**Supplementary Fig. 1**), leveraging the large cohort and the complementary capture of scRNA-seq and snRNA-seq. This yielded 54 fine-grained cell types (47 in scRNA-seq and 43 in snRNA-seq), most of which matched known identities and markers (**Supplementary Tables 6 & 9**), while also revealing populations that smaller studies typically miss.

Our approach of large-scale integration with accurate annotation provides four key advantages. First, it enables recovery of cell types that are essentially absent from previous heart single-cell atlases due to the inherent selective biases of most existing scRNA-seq technologies; for example, we detected rare basophils marked by *CLC* (0.1% in scRNA-seq; **Fig. 1g**). Second, it reveals rare cell types that have been underexplored in the heart; for example, the myeloid compartment harbors mature regulatory dendritic cells (mregDCs) marked by *LAMP3* (0.1% in scRNA-seq and 0.1% in snRNA-seq; **Fig. 1g, h**), a recently described dendritic-cell subset that promotes regulatory T (Treg) cell-mediated immune suppression in cancer^45^ and whose roles in the heart remain unclear. Third, it uncovers biologically interpretable states described in literature but underrepresented in cardiac single-cell data. Specifically, within the endothelial cells we identified a *DKK2*-expressing subset consistent with an angiogenic program, hereafter referred to as angiogenesis-associated endothelial cells (angiogenesis ECs) (**Fig. 1i**). *DKK2*-expressing endothelial cells were previously shown to promote angiogenesis in rodents and humans, but their role in the heart has not been well explored^46^. Fourth, with the recovery of such fine-grained subsets, our dataset expands on current cardiac cell atlases and can inform future investigations into their cellular functions and contributions to heart biology and disease. As we computed the DEGs in both the scRNA-seq and snRNA-seq data, the strong concordance between them indicates the robustness of our integrative approach (**Supplementary Fig. 2**) and thus gives confidence in the atlas and our results.

### Phenotype association analysis reveals the anti-apoptotic function of NRG1-expressing endocardial cells in cardiovascular diseases

With detailed cell-type annotations, we next examined how cell-type proportions correlate with sample phenotypes in the metadata, including age, sex, disease, and region. To create comparisons with sufficient sample size per category, we defined ten different phenotype comparisons (Old vs. Young, Female vs. Male, Disease vs. Normal, heart failure (HF) vs. Normal, dilated cardiomyopathy (DCM) vs. Normal, arrhythmogenic cardiomyopathy (ACM) vs. Normal, cardiac allograft vasculopathy (CAV) vs. Normal, Ventricle vs. Atrium, Left vs. Right, and AP vs. Ventricle). For each sample, we calculated the proportion of each cell type at three levels: broad cell type percentages, fine cell type percentages, and fine cell type percentages within their corresponding broad cell types. We then applied ANOVA to associate these percentages with phenotypic conditions, thereby identifying cell types associated with specific phenotypes (**Fig. 2, Supplementary Table 11**). For example, we found aging to be strongly associated with population changes in perivascular/endothelial cells and fibroblasts. Sex-based comparisons showed fewer associated cell types than other phenotypes, but CD56^+^ NK cells were significantly more abundant in females. In contrast, the disease phenotypes had the largest number of associated cell types, with different diseases sharing several overlapping cell types. Among these, CAV-related cell types showed the strongest associations and were primarily immune populations, consistent with the chronic alloimmune inflammation and vascular injury in CAV that broadly activate innate and adaptive immune cells. For heart regions, the most significant differences were observed between the ventricle and atrium, where atrial cardiomyocytes (aCMs) were exclusively found in the atrium and ventricular cardiomyocytes (vCMs) in the ventricle, supporting the reliability of our annotations. Additionally, pericytes showed differing proportions between the left and right regions, while endothelial cells displayed significant differences between the AP and the ventricle.

**Fig. 2:**
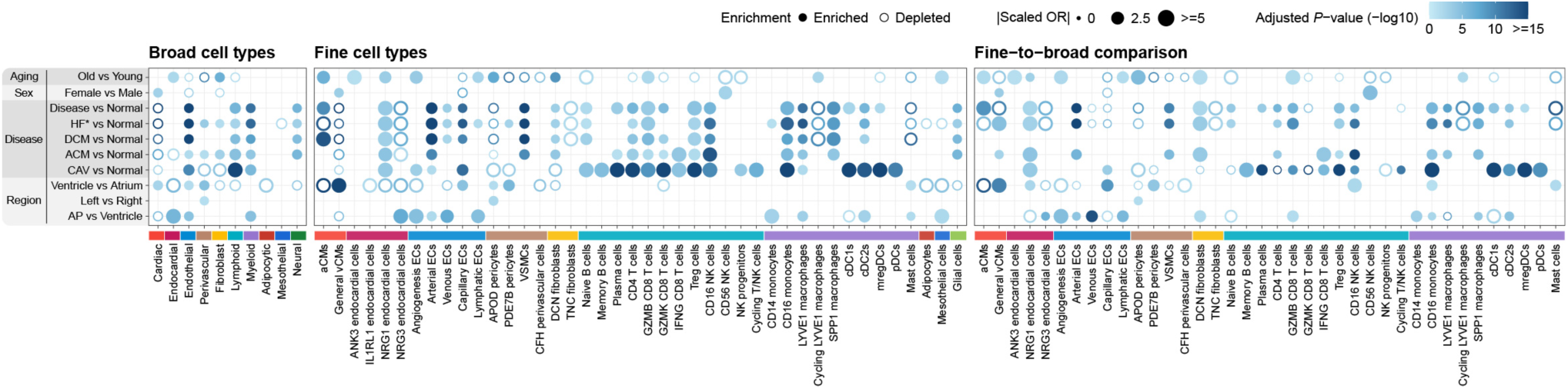
Association between cell type proportions and phenotypic features in snRNA-seq data. Dot plot showing the association results between the proportions of broad cell types and phenotypes (left), proportions of fine cell types and phenotypes (middle), and proportions of fine cell types relative to their respective broad cell types (right). Each dot represents a significant association (BH-adjusted *P*-value < 0.05). The color scale indicates the -log10 adjusted *P*-value, while the size of each dot corresponds to the absolute value of the scaled odds ratio (|scaled OR|). Filled and hollow circles denote enrichment and depletion of the corresponding cell type in the indicated phenotype, respectively. The comparison of “Old vs. Young” was performed using normal snRNA-seq samples only, and “HF*” includes heart failure samples as well as DCM and ACM samples diagnosed with heart failure.

Heart diseases often share common pathological features, such as cardiomyocyte loss and tissue remodeling. This raised the question of whether there exists a cell population that is consistently associated with multiple heart disease phenotypes. Interestingly, our analysis revealed a distinct subset characterized by high *NRG1* expression was consistently enriched across cardiovascular diseases in both snRNA-seq and scRNA-seq data (**Fig. 2**, **Fig. 3a**). Compared to controls, a higher proportion of the *NRG1* expressing endocardial cells were found in diseases, including DCM, restrictive cardiomyopathy (RCM)/left ventricular non-compaction (LVNC), ACM, and HF (**Fig. 3a**), with most of the DCM and ACM cases also presenting HF (**Supplementary Fig. 3a**). Notably, this enrichment was also observed in immune-driven CAV. Prior studies have shown that *NRG1* is an injury-induced growth factor that promotes cardiomyocyte proliferation and neovascularization, and is essential for trabecular development and heart failure progression^47,48^. *NRG1* also mitigates inflammation by suppressing tumor necrosis factor-alpha (TNFα) and interleukin-6 (IL-6) secretion^49^. To examine the association with other cardiovascular diseases, we analyzed an independent myocardial infarction (MI) dataset and found increased *NRG1*-expressing endocardial cells in the ischemic (IZ) and fibrotic (FZ) zones (**Fig. 3b; Supplementary Fig. 3b**). These results reinforce the role of *NRG1*-expressing endocardial cells across diverse cardiovascular conditions^47,50^. We further observed their elevated abundance in healthy aged hearts, suggesting potential involvement of *NRG1* in cardiac aging (**Fig. 2**).

**Fig. 3:**
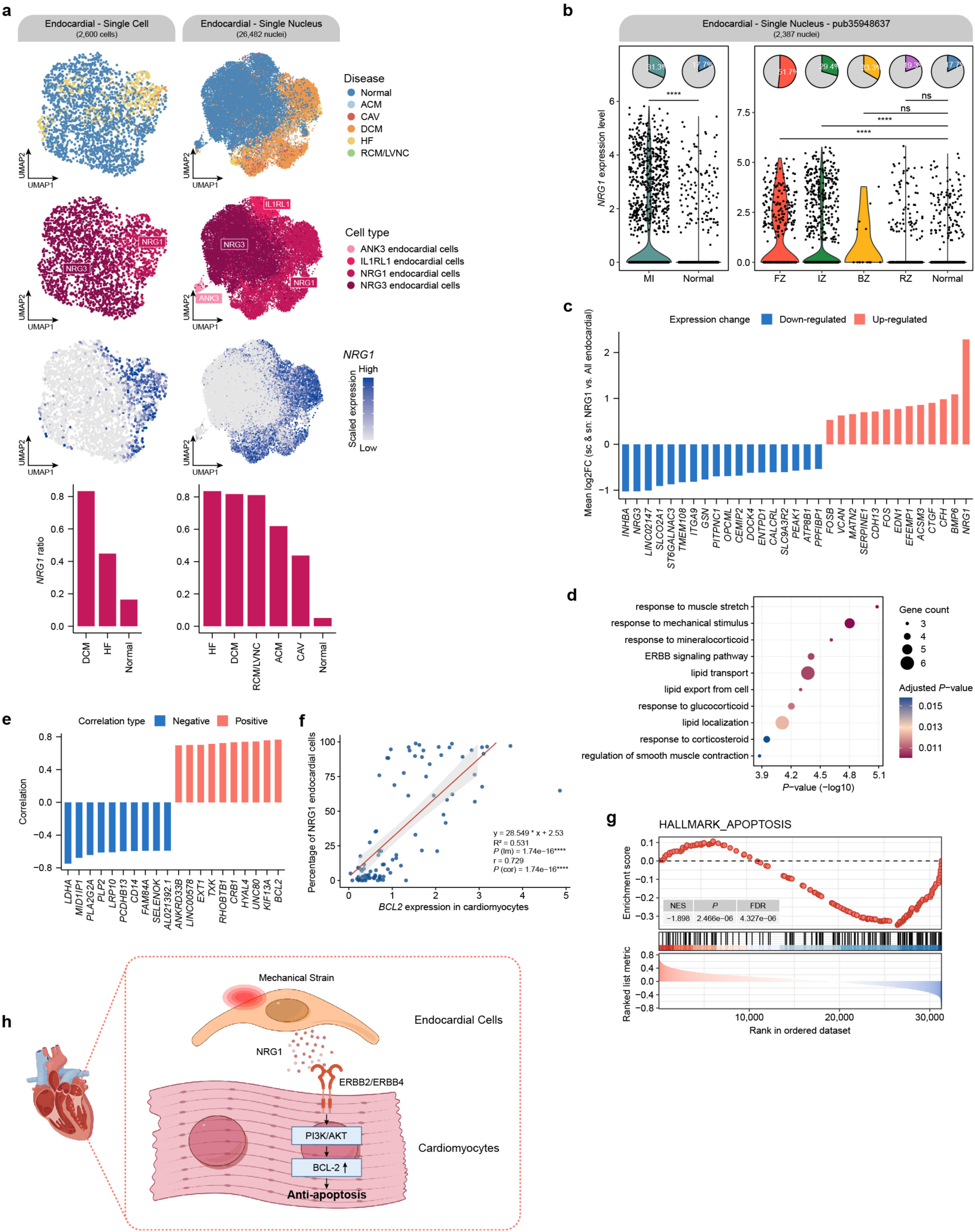
*NRG1*-expressing endocardial cells in pan-cardiac diseases. **a**, UMAP visualization of *NRG1*-expressing endocardial cells identified in the scRNA-seq and snRNA-seq data, highlighting specific endocardial subtypes, the *IL1RL1*, *ANK3*, *NRG1*, and *NRG3* subpopulations (2^nd^ row). Feature plots display the expression of *NRG1* (3^rd^), while a bar chart shows the proportion of *NRG1*-expressing endocardial cells across disease groups (4^th^ row). **b**, *NRG1* expression levels across myocardial infarction zones (FZ, fibrotic zone; IZ, ischemic zone; BZ, border zone; RZ, remote zone) and normal heart regions, demonstrating enrichment in fibrotic and injury zones. **c**, Differential expression analysis of *NRG1*-expressing endocardial cells by comparing against other endocardial cells to identify significantly up- and downregulated genes in scRNA-seq and snRNA-seq data. **d**, Gene set enrichment analysis (GSEA) of *NRG1*-expressing endocardial cells specific DEGs, revealing pathways associated with mechanical strain responses, ERBB signaling, and lipid transport. **e**, Identification of genes in cardiomyocytes whose expression levels strongly correlate with the proportion of *NRG1*-expressing endocardial cells within a sample. **f**, Correlation between the proportion of *NRG1*-expressing endocardial cells and *BCL2* expression levels in cardiomyocytes. **g**, GSEA results showing depletion of the APOPTOSIS hallmark gene set among genes correlated with the proportion of *NRG1*-expressing endocardial cells. **h**, Diagram illustrating the interaction between *NRG1*-expressing endocardial cells and cardiomyocytes, highlighting the anti-apoptotic effects mediated via ERBB2/ERBB4, PI3K/AKT, and BCL-2 signaling pathways (figure was created using Figdraw). Significance (two-sided Wilcoxon test) is indicated: nonsignificant (ns), \*\*\*\**P* < 0.0001.

In previous studies with bulk or small-scale single-cell datasets, they lacked the resolution and power to systematically characterize *NRG1*-expressing endocardial cells^20,24^. Here, we leveraged our assembled data to define their transcriptional program. We first identified DEGs with respect to the broader endocardial cell type in both scRNA-seq and snRNA-seq, and intersected the two DEG sets to reveal 31 shared genes (**Fig. 3c; Supplementary Tables 7 & 10**). Significant upregulation was observed for genes such as *NRG1*, *BMP6*, *CTGF*, and *EDN1* in the *NRG1*-expressing endocardial cells. Conversely, genes such as *INHBA*, *NRG3*, and *LINC02147* were notably downregulated. Gene Set Enrichment Analysis (GSEA) revealed that these DEGs were enriched in pathways related to “response to muscle stretch” and “response to mechanical stimulus,” aligning with previous studies and supporting their potential role in cardiac mechanotransduction and heart function (**Fig. 3d**)^51,52^.

Previous studies also revealed that *NRG1*-expressing endocardial cells interact with cardiomyocytes through ErbB receptors^52,53^. To explore their impact on cardiomyocytes, we correlated the proportion of *NRG1*-expressing endocardial cells with the gene expression profiles of cardiomyocytes across samples (**Fig. 3e, Supplementary Table 12**). Interestingly, *BCL2*, a key inhibitor of apoptosis, exhibited the highest positive correlation (**Fig. 3f**). Further analysis using GSEA revealed that the top positively correlated genes were negatively enriched in the apoptosis pathway, suggesting a potential anti-apoptotic role of these *NRG1*-expressing endocardial cells on cardiomyocytes and in regulating cardiomyocyte function (**Fig. 3g**). This is congruent with *NRG1*’s protective function in cardiovascular diseases.

In summary, our analysis identified *NRG1*-expressing endocardial cells as a disease-enriched, mechanosensitive niche that communicates with cardiomyocytes via ErbB receptors. This communication appears to confer anti-apoptotic protection in cardiomyocytes, as evidenced by the downstream *BCL2* upregulation and negative enrichment of apoptosis programs via PI3K/AKT. We posit that this is crucial in protecting cardiomyocytes during the early phase of ischemic or immune-mediated cardiac injury, which is congruent with the myocardium and priming regenerative responses previously attributed to NRG1/ErbB signaling (**Fig. 3h**)^53–55^. These findings are consistent with prior studies suggesting that endocardial *NRG1* is a therapeutic target for cytoprotection and repair in cardiovascular disease.

### Identification of aging-related signatures in different cell types

As one of primary risk factors for cardiovascular disease, aging impacts the heart at multiple levels, from functional to structural to cellular. The age altered heart environment thus provides the biological context in which pathological processes unfold. Here we followed up by investigating the aging-related transcriptional changes in normal cardiac samples. We first selected normal (non-diseased) samples from two large snRNA-seq datasets (EGAS00001006374 and ERP123138) for regression analysis^3,24^. By employing samples from a limited number of larger high-quality studies, we minimized batch effects that could confound the analysis. With the regression analysis, we identified age-correlated genes and visualized their changes across the ages. Interestingly, the average gene expression of these age-correlated genes remained relatively stable in most cell types until around 60 years of age, after which significant changes were observed, highlighting nonlinear changes in cardiac aging (**Fig. 4a**). This consistent pattern occurring across datasets suggests a general phenomenon rather than a project-specific observation and it aligns with epidemiological evidence that the risk of aging-related diseases rises nonlinearly over the human lifespan^56^.

**Fig. 4:**
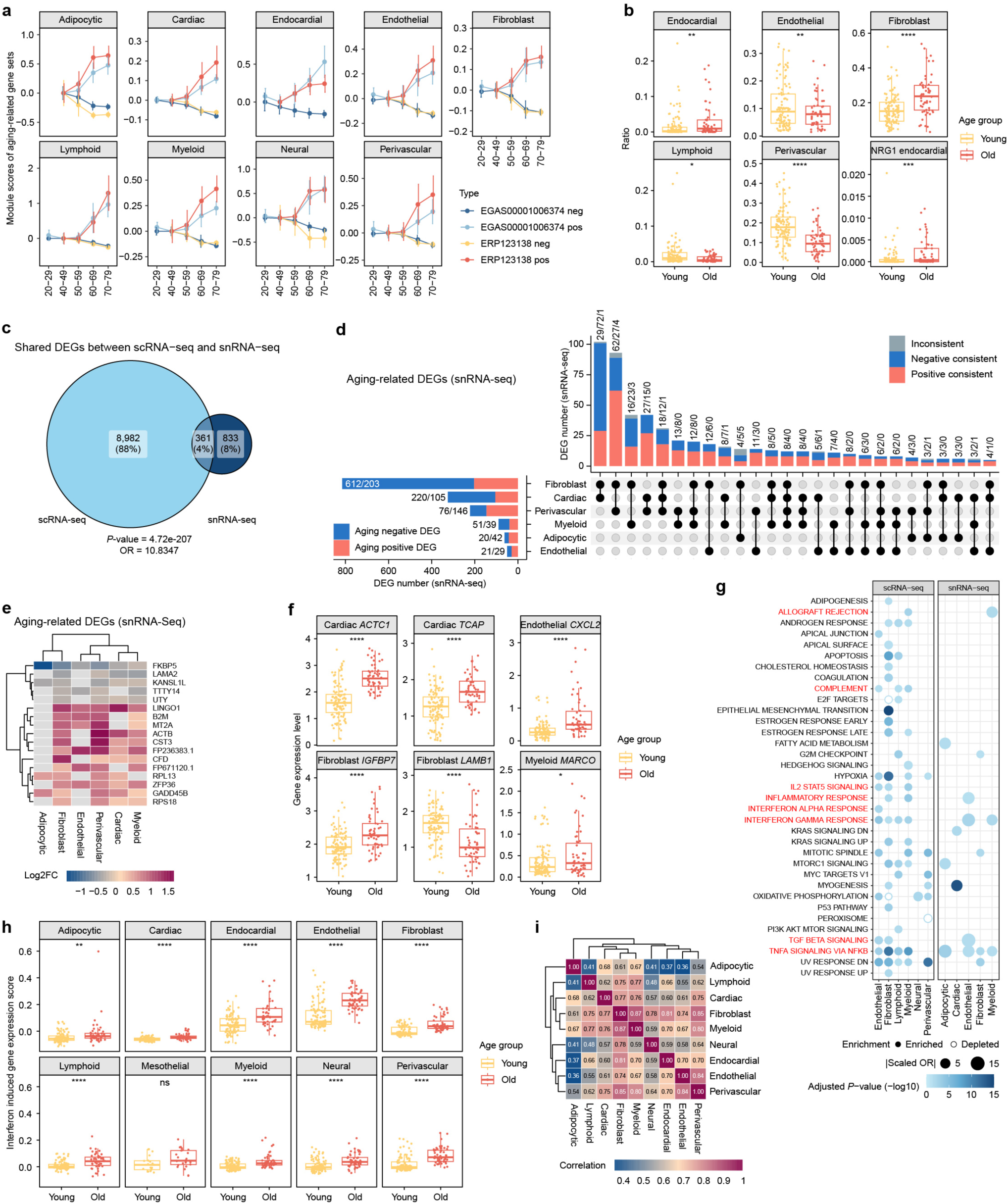
Comprehensive analysis of aging-related gene expression changes in various cell types. **a**, Module scores of aging-related gene sets found across different cell types in two selected datasets. **b**, Boxplot showing the proportions of six major cell types in the young and old samples. **c**, Venn diagram showing the overlap of aging-related DEGs in shared cell types between scRNA-seq and snRNA-seq data. **d**, Aging-related DEGs identified in the snRNA-seq data across major cell types. Left panel: Bar plot showing the total number of aging-related DEGs (positive and negative) per cell type. Numbers indicate upregulated (red) and downregulated (blue) DEGs in the old vs. young samples. Right panel: Upset plot illustrating overlaps of aging-related DEGs among cell types. Bars show the number of shared DEGs across combinations of cell types, with colors indicating directionality consistency: consistently positive (red), consistently negative (blue), or inconsistent (gray). **e**, Heatmap showing the expression patterns of aging-related DEGs shared across multiple cell types in snRNA-seq data. Rows represent genes; columns represent cell types. Color intensity indicates the log2 fold change (old vs. young), with red indicating upregulation and blue indicating downregulation. **f**, Representative examples of aging-related DEGs across different cardiac cell types. Boxplots show expression levels of selected genes in young (yellow) and old (red) individuals. **g**, ORA of aging-related DEGs across major cardiac cell types. For each cell type, dots represent significantly enriched Hallmark pathways (adjusted *P*-value < 0.05). Dot size corresponds to |scaled OR| (OR > 1 as enrichment; OR < 1 as −1/OR). Filled and hollow dots indicate enrichment and depletion, respectively. Color scale shows significance (-log10 adjusted *P*-value). Immune-related pathways are highlighted in red. Results are shown separately for scRNA-seq (left) and snRNA-seq (right). **h**, Boxplots showing the interferon-induced gene expression scores across ten cardiac cell types in young and old individuals. **i**, Heatmap showing inter-cell-type correlation of interferon response in snRNA-seq data. Significance (two-sided Wilcoxon test) is indicated: nonsignificant (ns), \**P* < 0.05, \*\**P* < 0.01, \*\*\**P* < 0.001, \*\*\*\**P* < 0.0001.

Based on this observation and to facilitate differential expression and other analyses, we categorized the samples as “young” (<60 years) or “old” (≥60 years). This binary classification ensured a substantial number of samples in each group, reducing batch effects and improving statistical power. We first compared the cell type composition between young and old hearts and found aging-related population shifts like increase in fibroblasts and decrease in perivascular cells (**Fig. 4b**). Notably, the directionality of these changes matched our earlier analysis with the whole atlas (**Fig. 2**). Here we also examined the *NRG1*-expressing endocardial subset and found the same trend of increasing proportion with age in healthy samples. This combined with our previously identified association between cardiovascular disease and increased abundance of *NRG1*-expressing endocardial cells, highlights potential connections between aging and cardiovascular disease and suggests a role for *NRG1* in the process by which aging predisposes the heart towards disease (**Fig. 4b**).

We next computed the aging-related DEGs among the broad cell types. Comparison between the old and young groups identified 9,343 and 1,581 aging-related DEGs from the scRNA-seq and snRNA-seq datasets, respectively (**Supplementary Tables 13 & 14**). Intersecting aging-related DEGs from both platforms yielded a small but highly significant overlap (4%, 361 genes; *P*-value = 4.72×10^-207^, OR = 10.83; **Fig. 4c**), indicating clear cross-platform concordance. In the snRNA-seq data, fibroblasts, cardiomyocytes, and perivascular cells showed the highest number of aging-related DEGs (**Fig. 4d**). In contrast, in the scRNA-seq data, myeloid cells, endothelial cells, and perivascular cells displayed the most aging-related DEGs (**Supplementary Fig. 4e**). These results emphasize the need for complementary methods to capture the full spectrum of aging-related gene expression changes. Although most DEGs were cell type-specific, shared DEGs were observed between cell type pairs, such as fibroblasts and cardiomyocytes, as well as fibroblasts and perivascular cells (**Fig. 4d**). These shared DEGs tended to change in the same direction, indicating similar aging effects across cell types. Additionally, aging-related negative DEGs outnumbered positive DEGs in both data modalities, suggesting a general downregulation of gene expression with aging.

Focusing on the shared aging-related DEGs, we identified 17 and 144 common aging-related DEGs in snRNA-seq (shared by at least four cell types) and scRNA-seq data (shared by at least five cell types), respectively (**Fig. 4e; Supplementary Fig. 4f**). Notably, these common genes are largely related to fundamental housekeeping functions affected during aging. For instance, we found mitochondrial-encoded genes (*MT-CYB*, *MT-CO1*, *MT-CO3*, and *MT-ND1*) to be consistently upregulated across cell types. This increase likely reflects a compensatory response to mitochondrial dysfunction, a hallmark of aging associated with elevated reactive oxygen species (ROS)^57^. Similarly, metallothionein genes (*MT1M*, *MT1X*, and *MT2A*) were upregulated across multiple cell types, underscoring their role in oxidative stress mitigation and metal ion regulation during aging^58^. Ribosomal protein genes, critical for protein synthesis and stress response, also increased with age, likely reflecting an adaptive response to proteostasis imbalance caused by accumulated misfolded or damaged proteins^59^.

Examining the cell type-specific aging-related DEGs, we found genes that are closely linked to the specialized functions of each cell type (**Fig. 4f**). In cardiac cells, *ACTC1* (alpha cardiac actin 1) and *TCAP* (titin-cap protein) exhibited significantly higher expression levels in older individuals compared to younger ones. These genes are essential for cardiac contractility and sarcomere integrity, indicating potential aging-related remodeling in cardiac muscle cells to adapt to functional demands^60,61^. In endothelial cells, *CXCL2*, a pro-inflammatory cytokine, showed a marked increase in expression with aging. This upregulation suggests a heightened inflammatory response, which may contribute to endothelial dysfunction, a hallmark of vascular inflammaging^62^. In fibroblasts, *IGFBP7* (insulin-like growth factor binding protein 7) was significantly upregulated, while *LAMB1* (laminin subunit beta 1) was downregulated in older individuals. These genes are involved in extracellular matrix remodeling and cell adhesion, highlighting their contributions to age-associated fibrosis and tissue stiffening^63,64^. Similarly, *MARCO* (macrophage receptor with collagenous structure) exhibited a modest but significant increase in expression in myeloid cells of the older group. Known for its role in immune response, this suggests an aging-related shift in immune cell functionality, potentially reflecting chronic low-grade inflammation often observed during aging^65^.

We also identified aging-related DEGs at the fine cell type level (**Supplementary Fig. 4a; Supplementary Table 15**). One notable gene is *NPPA*, which encodes the atrial natriuretic peptide, and it exhibits higher gene expression levels in the atrial cardiomyocytes of aged hearts (**Supplementary Fig. 4c**). *NPPA* has been recognized as a significant marker of cardiac aging in prior studies, as exemplified by its observed upregulation in aged human induced pluripotent stem cell (iPSC)-derived cardiomyocytes^66^. We then conducted a thorough examination of *NPPA* expression with respect to aging by checking its expression across the different heart regions and age groups. We found that elevated *NPPA* expression primarily occurs in the atrium and it can begin as early as 40 years of age (**Supplementary Fig. 4c**). This suggests that while large-scale gene expression changes tend to occur at a later age (>60 years), some smaller-scale changes have already begun a decade or more earlier. Taken together, our DEG analysis of large-scale, regionally resolved human datasets establishes atrial *NPPA* upregulation as a robust feature of human cardiac aging.

### Widespread interferon-related inflammation across diverse cell types in cardiac aging

To elucidate the functional implications of the aging-related DEGs that we identified, we performed over-representation analysis (ORA) with Molecular Signatures Database (MSigDB) Hallmark gene sets and reported the effect sizes as ORs (**Fig. 4g**). Several immune and inflammatory pathways (highlighted in red), including allograft rejection, complement, IL2/STAT5 signaling, inflammatory response, interferon alpha response, interferon gamma response, TGF beta signaling, and TNF-α signaling via NF-κB, are significantly enriched across many non-cardiomyocyte populations like endothelial cells, fibroblasts, and myeloid cells. This expression pattern reveals a predominantly inflammatory phenotype centered on TNF and IFN signaling during cardiac aging. Such an elevated interferon response is frequently associated with the activation of the cyclic GMP–AMP synthase (cGAS)–stimulator of interferon genes (STING) pathway, a recognized hallmark of aging^67,68^.

To investigate the IFN responses during aging in different cell types, we computed an IFN score using all IFN response genes (**Supplementary Fig. 4d**). Here we found significantly higher IFN response scores in endothelial cells, mesothelial cells, and perivascular cells compared to other cell types. When comparing between the old and young, we observed increased IFN response scores across most cell types in the aged samples. In particular, endothelial cells, endocardial cells, perivascular cells, and fibroblasts exhibited the most significant changes (**Fig. 4h**). These findings are congruent with the role of the IFN response in driving age-associated changes in the cellular microenvironment, particularly in vascular and fibroblast populations^69^.

To explore whether these responses were coordinated across cell types, we calculated pairwise correlations of IFN scores between cell types. As shown in **Fig. 4i**, endothelial, endocardial, perivascular cells, and fibroblasts display strong correlation between their IFN scores (r > 0.8), suggesting synchronized activation. Notably, these same cell types also display strong intercellular communications in our PopComm analysis (refer to next section). This pattern raises the possibility that cGAS–STING signaling may mediate a cascade of IFN responses across multiple cell types, potentially through paracrine mechanisms. Such coordinated activation can contribute to widespread inflammatory remodeling in the aging heart.

### Cell-level convergence toward aging-associated states in cardiac aging

Although comparisons at the cell type level effectively capture overall age-associated differences, they may mask gradual or continuous shifts in cellular states that occur within individual lineages during aging. Building on cell type–level analyses (**Fig. 4**), which revealed broad transcriptional differences between young and old cardiac samples, we next examined whether aging also introduces within-cell-type heterogeneity at the single-cell level.

We first quantified an aging signature for each cell by calculating the difference between the mean expression of positively and negatively regulated aging-related DEGs within its cell type. This approach provides a quantitative framework to assess continuous aging trends within each lineage. When visualizing the aging signature scores on Uniform Manifold Approximation and Projection (UMAP) embeddings (**Fig. 5a, b**), cells with high scores showed non-random localization patterns across the plots, which differed among cell types. Although UMAP-based visualization provides only a qualitative representation of transcriptional similarity and does not fully preserve the global data structure, these patterns nonetheless suggest that aging-associated gene programs are preferentially enriched within specific regions of the transcriptional landscape for each lineage. Together, these distributions indicate that aging does not act uniformly within a lineage but instead remodels molecular programs in distinct ways across cell types. Aging likely activates shared processes such as stress responses, metabolic remodeling, and inflammatory signaling. As a result, subsets of cells begin to express overlapping aging-related gene programs, giving rise to a convergence of cellular states within each lineage that reflects a common aging-associated transcriptional landscape.

**Fig. 5:**
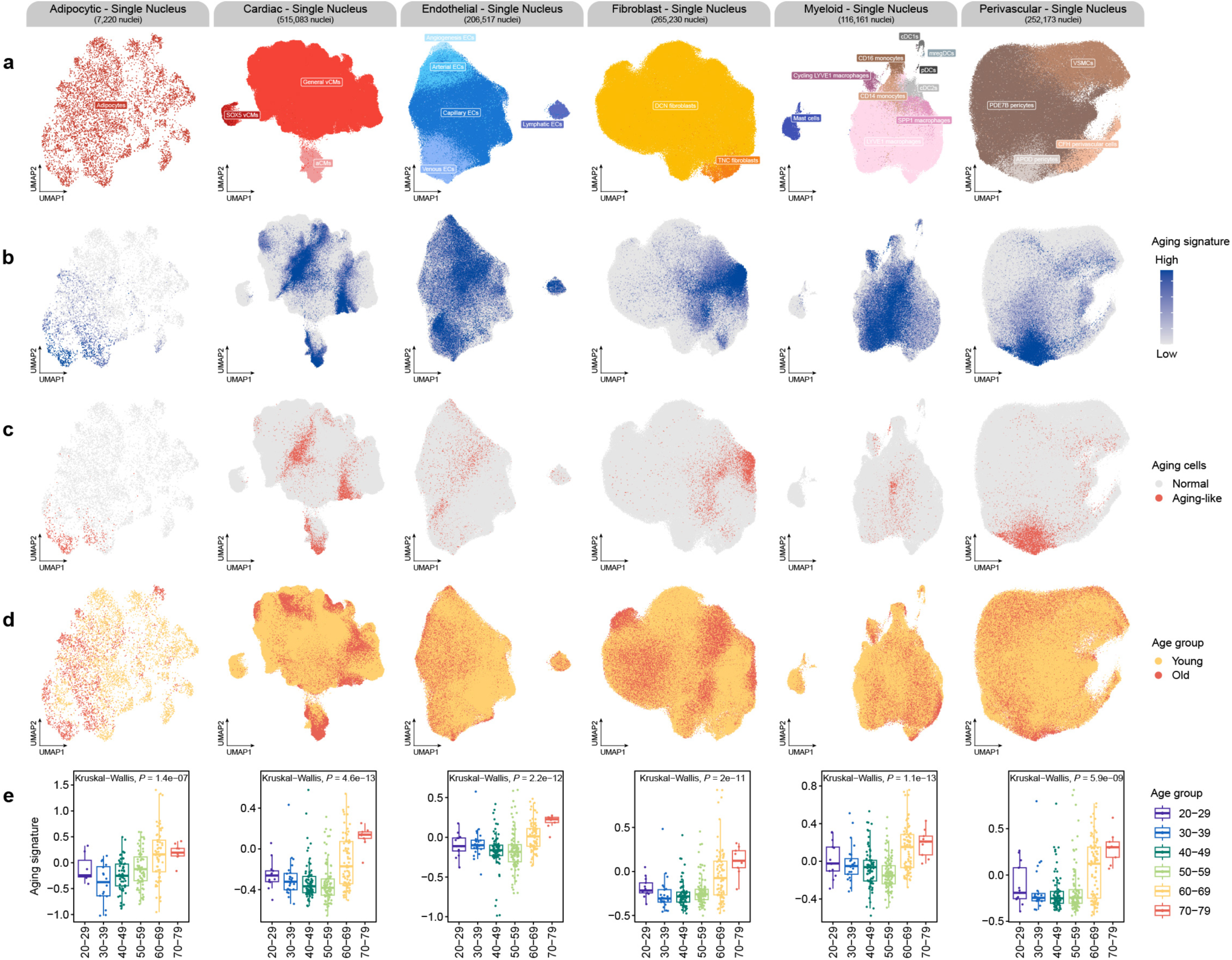
Aging signature patterns and senescent cell distribution across cardiac cell populations. **a**, UMAP visualization of subset annotations for six major cardiac cell types. **b**, Visualization of aging signature scores across six major cardiac cell types. **c**, UMAP plots show the localization of aging-like cells (red). Aging status was defined based on transcriptional signatures, and these cells appear enriched in specific cell types, particularly in fibroblasts, endothelial, and perivascular cells. **d**, Visualization of age group (young and old) distribution across six major cell types. **e**, Aging signature scores across different age groups for six major cell types.

Based on this analysis, we classified aging-like cells of each lineage using a cutoff at the 95^th^ percentile of the aging signature score (**Fig. 5c**). Interestingly, within adipocytes, fibroblasts, and perivascular cells, the aging-like cells formed distinct clusters within their respective cell types. For instance, the aging perivascular cells were almost entirely localized to *APOD* pericytes. For the cardiomyocytes, three distinct groups of aging-like cells were identified, potentially representing different subsets of cardiomyocytes influenced by aging in diverse ways. In contrast, aging-like cells in the endothelial and myeloid cells were more diffusely distributed, suggesting a broader, less localized effect of aging on these cell types. This heterogeneity in the distribution of aging-like cells highlights the cell type-specific responses to aging, with some cell types showing a more pronounced convergence toward aging-associated states than others.

We further compared the distribution of aging-like cells with age information in the metadata (**Fig. 5d**). Adipocytes, cardiomyocytes, and the *APOD* pericyte subset showed relatively good alignment between aging signature scores and age groups. Fibroblasts displayed a more moderate correspondence, with aging-like nuclei enriched in older donors but also present in young hearts. By contrast, endothelial and myeloid cells showed only limited agreement, as high aging scores were observed across donors regardless of age. These observations suggest that gene-expression-based aging signatures can reveal aging-related cellular states that are not fully captured by chronological age alone.

Additionally, we calculated the average aging signature score of each cell type in each sample and binned them by age. Consistent with our findings in the two datasets (EGAS00001006374 and ERP123138), the aging signature changed significantly after 60 years of age (**Fig. 5e**), reinforcing our conclusion on a critical aging breakpoint in the heart.

### PopComm: inference and quantification of ligand-receptor interactions from population-scale single-cell data

Proper functioning of the heart depends not only on the individual roles played by each cell type but also on the cell-cell communications that give rise to the emergent properties of the heart. Therefore, dissecting cell-cell interactions in normal physiological states, as well as their alterations during aging and pathological conditions, is an important aspect needed for understanding the mechanisms underlying normal heart functions and disease progression. With whole-transcriptome single-cell data, we can perform large-scale screening to infer cell-cell interactions by assessing the gene expression patterns of ligand and receptor genes. The feasibility of this approach has been demonstrated by methods like CellPhoneDB^40^ and CellChat^41^. However, these methods are primarily designed for smaller datasets and focus mainly on inferring cell-cell interactions while lacking capabilities for sample-level quantification and phenotype association analysis. As our heart data is composed of 79 scRNA-seq and 357 snRNA-seq samples, existing methods are insufficient to account for such complexity, underscoring the need for a more robust sample-level analysis approach.

To address these limitations, we developed PopComm, a toolkit designed for the inference and quantification of ligand-receptor interactions from population-scale single-cell data (**Fig. 6**). The core hypothesis of PopComm is that if a ligand-receptor pair mediates communication between two cell types, the expression levels of the ligand in the sender cell type and receptor in the receiver cell type should demonstrate strong positive correlation across diverse individuals. PopComm filters for statistically significant ligand-receptor pairs and quantify the communication intensities at the sample-level. PopComm also includes multiple visualization functions and is available in both R and Python versions. Detailed descriptions of each analytical step and the mathematical formulations underlying PopComm can be found in the Methods section and Supplementary Materials.

**Fig. 6:**
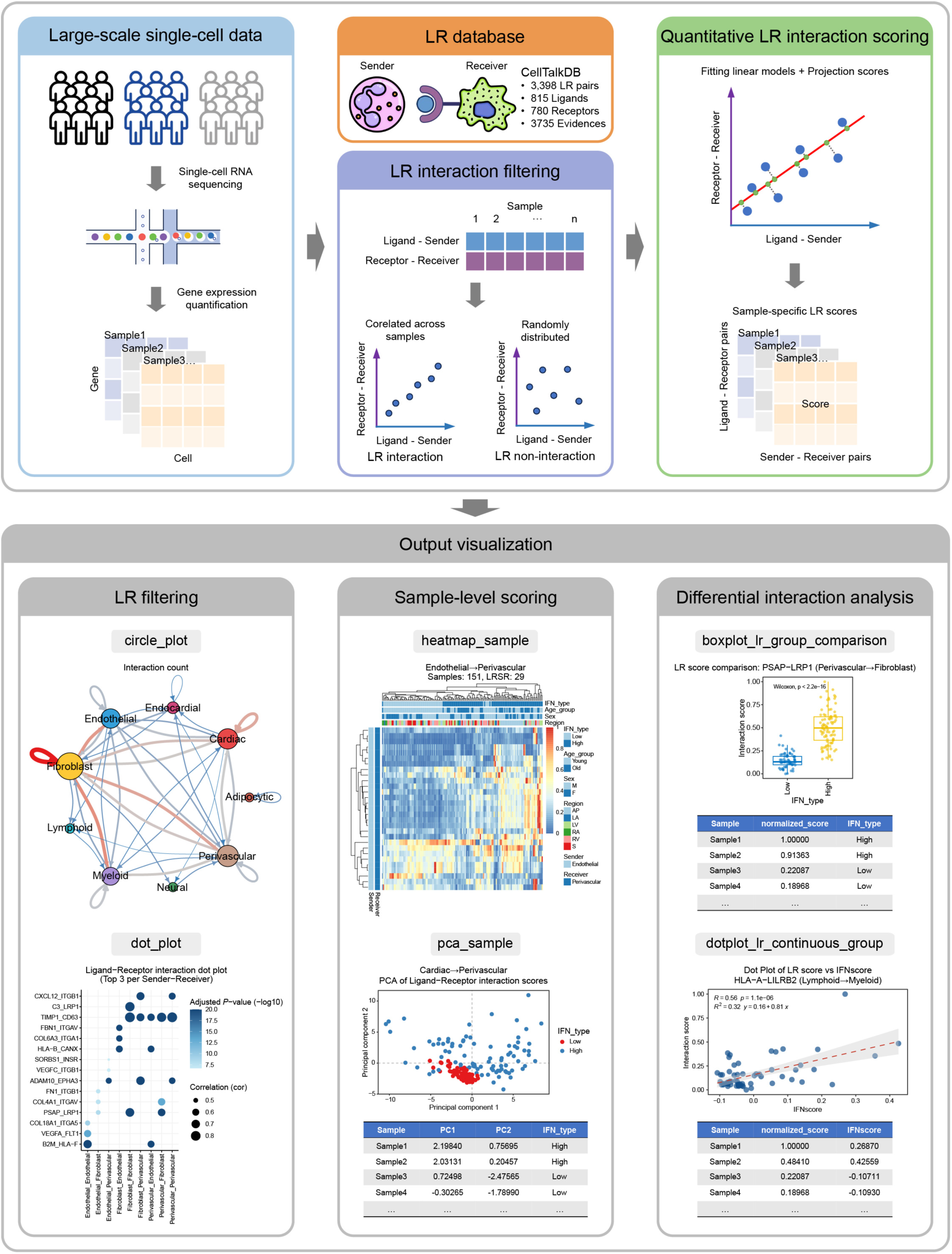
Overview of PopComm. Top panel: An overview of the statistical framework for inferring two cell-type-specific ligand-receptor pairs and quantifying sample-level cell-cell communication from large-scale single-cell transcriptomics data, along with a summary of the utilized databases. Bottom panel: Illustration of the six output visualization modules, organized into three levels: ligand-receptor filtering (LR filtering), sample-level scoring (Sample-level Scoring), and differential interaction analysis (Differential interaction analysis).

Due to the differences in cell type composition and gene expression profiles, we applied PopComm separately to our single-cell and single-nucleus heart atlases. Initial analysis identified a total of 1,106 and 2,507 ligand-receptor-sender-receiver (LRSR) pairs in the scRNA-seq and snRNA-seq datasets, respectively. Among these, 325 pairs were shared between the two modalities, showing significant overlap (*P*-value = 0, OR = 44.14; **Fig. 7a; Supplementary Tables 17 & 18**). In the scRNA-seq data, interactions between endothelial cells and pericytes were the most prominent, along with strong self-interactions within each of these cell types. In contrast, the snRNA-seq data revealed fibroblasts as a significant hub with extensive interactions with endothelial cells, pericytes, myeloid cells, and cardiomyocytes (**Fig. 7b**). Strong self-interactions were also observed in the fibroblasts as well as cardiomyocytes in the snRNA-seq data. We further observed that the number of LR pairs identified per cell type was highly correlated with the number of individuals in which that cell type was detected (**Supplementary Fig. 5a**). After correcting for sample size effects, fibroblasts emerged as having the highest number of interactions in both scRNA-seq and snRNA-seq datasets, whereas lymphoid cells consistently exhibited fewer interactions.

**Fig. 7:**
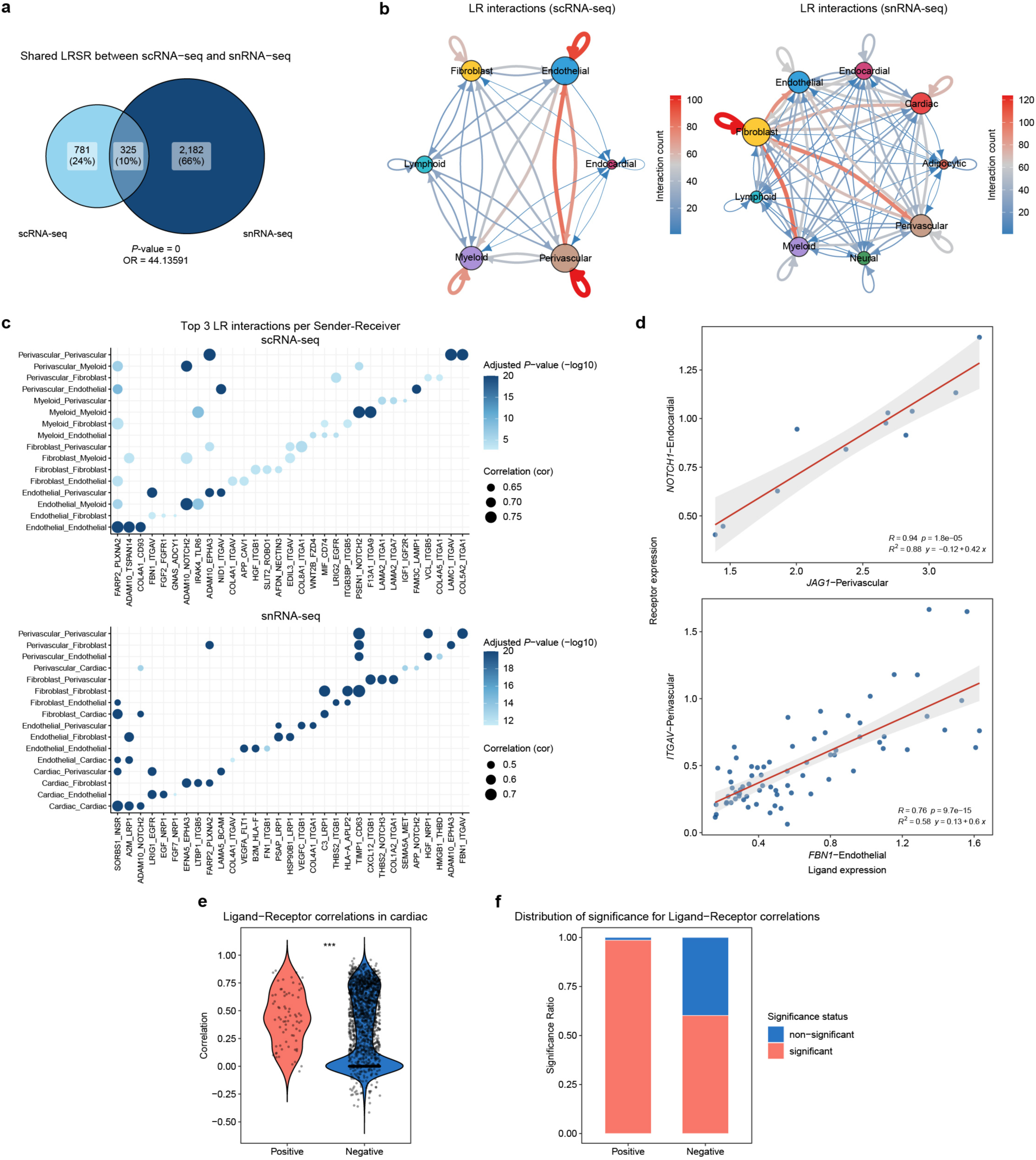
PopComm reveals consistent and spatially validated ligand–receptor interactions across single-cell and single-nucleus transcriptomes. **a**, Venn diagram showing the overlap of ligand–receptor-sender-receiver (LRSR) pairs identified from scRNA-seq and snRNA-seq datasets. **b**, Circle plots showing the number of significant ligand–receptor (LR) interactions among major cell types in the scRNA-seq (left) and snRNA-seq (right) datasets. **c**, Dot plots showing the top 3 LR interactions per sender–receiver pair. Dot size indicates the correlation coefficient across individuals, and color intensity represents the adjusted *P*-value of the interaction. Top: interactions involving Fibroblast, Endothelial, Perivascular, and Myeloid cells (scRNA-seq). Bottom: interactions involving Fibroblast, Endothelial, Perivascular, and Cardiac cells (snRNA-seq). **d**, Scatter plots validating two representative LR interactions by correlating ligand and receptor expression across samples. **e**, Boxplot showing the spatial correlation of LR gene expression in cardiomyocyte-enriched spots in a 10x Genomics Visium spatial transcriptomics dataset. **f**, Bar plot showing the proportion of LR pairs with statistically significant spatial correlations (*P*-value < 0.01) in the “Positive” (PopComm-predicted) and “Negative” (non-predicted) groups. Significance (two-sided Wilcoxon test) is indicated: \*\*\**P* < 0.001.

At the level of individual LR pairs, PopComm successfully identified well-established interactions (**Fig. 7c; Supplementary Fig. 5b, c**). For example, PopComm recovered the vascular signaling such as JAG1–NOTCH1 between perivascular and endothelial cells^70^, as well as matrix–integrin signaling exemplified by FBN1–ITGAV between endothelial and perivascular cells^71^ (**Fig. 7d**). These results illustrate PopComm’s ability to capture both known and novel communication axes, thus providing a systematic framework for mapping cell–cell interactions in large-scale dataset such as our heart atlas.

We then employed a spatial transcriptomics (Visium) dataset to validate the LR interactions predicted by PopComm. Given the dominant presence of cardiomyocytes in the spatial data, our validation focused on cardiomyocyte self-interactions. For each LR pair, we calculated the spatial correlation of ligand and receptor expression across spots with a high cardiomyocyte signal (**Fig. 7e**). We next compared the PopComm-predicted pairs (“Positive”) with pairs not predicted but expressed in the data (“Negative”). Strikingly, the PopComm-predicted LR pairs showed significantly higher correlation values and a larger proportion of statistically significant interactions compared with the background set (*P*-value < 0.01; **Fig. 7f**). These findings support the reliability of PopComm in identifying spatially coordinated ligand–receptor interactions.

### Disease status and interferon responsiveness shape cell-to-cell interactions in aging hearts

PopComm outputs sample-level LR interaction scores which allows us to stratify patients based on their L-R interaction patterns. Here, we associated L-R interaction score with sample metadata. In the snRNA-seq dataset, unsupervised clustering of the LR scores showed coherent groupings of disease labels, implying that cardiac disease status was the dominant factor associated with the cell–cell interactions, whereas other factors such as heart region did not have labels forming coherent groups (**Fig. 8a**).

**Fig. 8:**
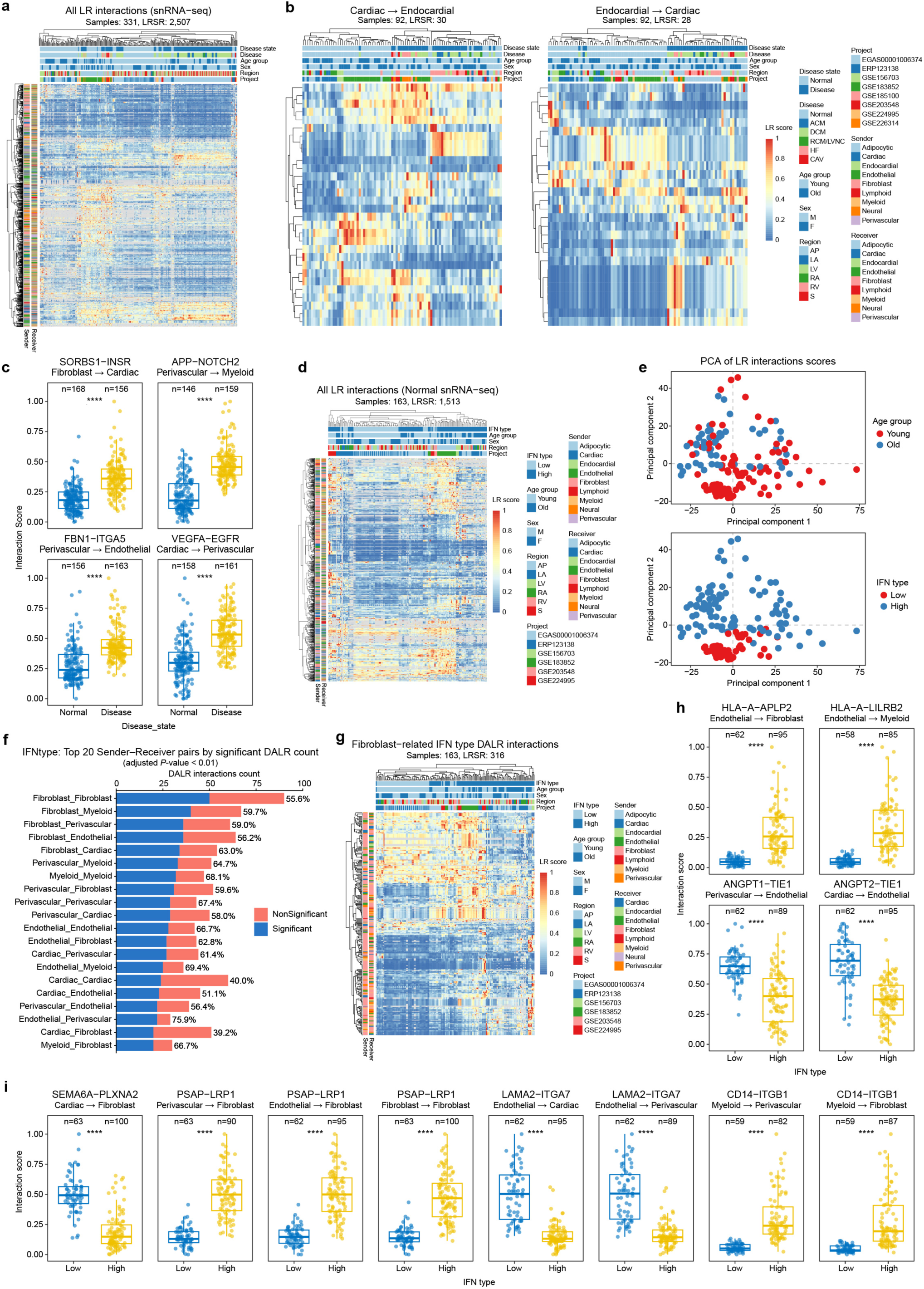
PopComm uncovers differential ligand–receptor interactions linked to aging and IFN-related signaling. **a**, Heatmap showing sample-level scores of 2,507 ligand–receptor (LR) interactions inferred by PopComm from the snRNA-seq data with 331 samples. Rows represent LRSRs and columns represent individual samples, with annotation by cell type proportions and donor metadata. **b**, Heatmaps showing directional LRSR activity from cardiomyocytes to endocardial cells (left) and from endocardial to cardiomyocytes (right). **c**, Boxplots showing representative LR interaction scores associated with heart disease status. **d**, Heatmap of LRSR scores restricted to normal snRNA-seq samples (n = 163), covering 1,573 interactions. **e**, PCA of sample-level LR interaction scores inferred by PopComm. Each dot represents a sample. The left panel is colored by age group (old vs. young), and the right panel is colored by interferon status (IFN type) (high vs. low). **f**, Bar plot showing the top 20 sender–receiver cell-type pairs ranked by the number of differentially active LRSRs (DALRs) associated with interferon status. **g**, Heatmap showing fibroblast-related LRSRs significantly associated with interferon status across 158 samples. **h-i**, Boxplots of representative interferon–associated LRSRs across sender–receiver pairs. Significance (two-sided Wilcoxon test) is indicated: \*\*\*\**P* < 0.0001.

Interestingly, we also found that different sender–receiver cell type pairs exhibited distinct associations with disease. Among all sender-receiver interaction pairs, interactions involving cardiomyocytes and endocardial cells showed the significant differences between disease and non-disease samples in the unsupervised clustering analysis (**Fig. 8b**). We further identified differentially active ligand–receptor (DALR) pairs between disease and non-disease samples for each sender–receiver cell type pair (**Supplementary Table 20**). Notable examples include the SORBS1–INSR interaction between fibroblasts and cardiac cells, which is involved in metabolic regulation under heart stress conditions. Another is the APP–NOTCH2 interaction between perivascular and myeloid cells that plays a role in modulating inflammation within the cardiac microenvironment. The FBN1–ITGA5 interaction between perivascular and endothelial cells is responsible for vascular remodeling and may contribute to fibrosis and tissue repair in the heart. Finally, the VEGFA–EGFR interaction between cardiac and perivascular cells influences angiogenesis and vascular remodeling in ischemic heart conditions. These interactions highlight how disease status can impact cell communication, regulating inflammation, fibrosis, and vascular repair (**Fig. 8c**).

Unsupervised clustering based on the LR scores from all healthy control samples in the snRNA-seq dataset suggested that age group might be a dominant factor in control samples influencing cell–cell interactions (**Fig. 8d; Supplementary Table 19**). Furthermore, we showed earlier that the expression of IFN response genes was highly associated with aging. Therefore, we stratified the control samples into two groups based on their expression levels of IFN response genes. Interestingly, the IFN response status showed a stronger association with global LR scores than chronological age group. Principal component analysis (PCA) analysis of the LR score matrix further supported this observation, with principal components (PC1 and PC2) being more strongly correlated with IFN response status than with age group (**Fig. 8e**). These results suggest that IFN signaling may act as a key mediator of aging-related changes in cell–cell communication.

We next identified DALR pairs between the IFN-high and IFN-low groups. Among all cell types, fibroblasts exhibited the highest number of DALRs as either sender or receiver (**Fig. 8f; Supplementary Table 21**). Unsupervised clustering based on the LR scores involving fibroblasts in either sender or receiver roles resulted in a clearer separation between the IFN groups as compared to clustering using LR scores from all cell types (**Fig. 8d, 8g**). This highlights the central role of fibroblasts in IFN-associated cell–cell communication changes.

We also note that several of the most prominent DALRs showed strong biological relevance. For instance, the HLA-A–APLP2 and HLA-A–LILRB2 interacting pairs were significantly upregulated in IFN-high samples, which is consistent with enhanced antigen presentation and immunoregulatory signaling. Conversely, the ANGPT1/2–TIE1 interactions, key for maintaining endothelial stability, was downregulated, possibly reflecting increased endothelial activation or permeability in IFN-rich conditions (**Fig. 8h**). We also found the SEMA6A–PLXNA2 interaction between cardiomyocytes and fibroblasts to be reduced in both aging and IFN-high states, marking it as a potential signature of pathological tissue remodeling. In contrast, PSAP–LRP1 interactions, originating from multiple lineages and targeting fibroblasts, were strongly upregulated, with high ligand–receptor co-expression, suggesting the activation of matrix turnover or immune clearance pathways. Additional IFN-sensitive interactions, such as LAMA2–ITGA7 and CD14–ITGB1, further support the idea that the IFN response promotes remodeling of the extracellular matrix and immune–stromal crosstalk (**Fig. 8i**).

Together, these findings indicate a tight coupling between IFN signaling and cell–cell communication, particularly in pathways related to immune regulation, vascular remodeling, and stromal activation during cardiac aging.

## Discussion

In this study, we integrated single-cell and single-nucleus data from 12 studies to construct a comprehensive cardiac atlas comprising 355,762 cells and 1,436,719 nuclei. The corresponding sample metadata were meticulously collected and curated, resulting in 36 metadata fields. Comparative analysis revealed significant differences in cell capture efficiency and gene expression profiles between scRNA-seq and snRNA-seq platforms, underscoring the necessity of integrating both technologies to achieve comprehensive and consistent cell type annotation across studies. We employed a hierarchical annotation approach, which facilitated the identification of 10 broad cell types and 54 fine cell types. One of the key innovations in our study is the development of PopComm, a novel tool designed to infer ligand-receptor interactions from population-scale single-cell data and quantify interaction strength at the individual sample level. Using PopComm, we found that IFN response states influence cell–cell communication during cardiac aging, offering new insights into the mechanisms underlying aging-related cardiac dysfunction. Additionally, we identified *NRG1*-expressing endocardial cells as being enriched in several cardiac diseases, enhancing our understanding of the molecular basis of heart disease. This integration approach and the development of PopComm address the limitations of previous single-cell atlases, which often lacked sufficient statistical power for phenotype association analysis, especially in the context of cardiovascular diseases and aging. These findings provide a valuable resource for future investigations into the cellular and molecular mechanisms of cardiac health and disease.

Our study also reveals the existence of rare cell types, such as adipocytes (which are likely to be derived primarily from epicardial fat, given the diverse anatomic sampling in our data), and poorly defined populations, including angiogenesis-associated endothelial cells (angiogenesis ECs), mature regulatory dendritic cells (mregDCs), and basophils. These populations exemplify cell types that were previously underrepresented or undefined in traditional cardiac atlases. Leveraging large-scale datasets with advanced integration, our atlas robustly detects rare lineages, systematically recovers underexplored cell types, and refines cell-type phenotypes, thus providing an actionable reference to connect cellular heterogeneity with phenotypic variation and disease. Further research into the biological functions of these cell types can offer valuable insights for identifying new therapeutic targets for heart diseases.

We grouped the curated phenotypes into four broad categories: age, disease, gender, and heart region, each with several subcategories. Cell type proportions within each sample were then associated with these phenotypes to identify phenotype-associated cell types. Focusing on disease, we discovered that *NRG1*-expressing endocardial cells were enriched across all disease types in our data, in both single-cell and single-nucleus modalities. Validation with external datasets confirmed this finding, showing that *NRG1*-expressing endocardial cells are enriched in the fibrotic zone of myocardial infarction, suggesting their potential role in fibrosis. Further analysis revealed that the marker genes of *NRG1*-expressing endocardial cells are significantly associated with mechanical stimuli, which may explain their activation in diseased states^52,72^. Additionally, we found that these cells interact with cardiomyocytes and activate anti-apoptotic pathways in the latter. This interaction may enhance cardiomyocyte survival in a diseased microenvironment, which could in turn influence tissue remodeling processes. These findings suggest that *NRG1*-expressing endocardial cells could be potential therapeutic targets for mitigating cardiac fibrosis in various disease contexts^73,74^. Additionally, some macrophages were found to be enriched in multiple cardiovascular disease phenotypes, further supporting their role in inflammation and tissue remodeling^75^.

Leveraging our large dataset, we conducted a comprehensive analysis of cardiac aging. Interestingly, our findings reveal that cardiac aging is not a linear process, with significant alterations occurring after the age of 60. Differential expression analysis demonstrated that aging-related DEGs in each cell type are closely linked to cell type-specific functions. Additionally, some aging-related DEGs are associated with shared pathways, such as “MYC Targets”, “Oxidative Phosphorylation”, Mitotic Spindle” and “UV Response DN”, among others. Notably, we observed a significant increase in IFN signatures, such as “Interferon Alpha Response” and “Interferon Gamma Response,” across almost all cell types, suggesting that the activation of interferon signaling may play a significant role in the aging process across multiple cell types. These findings provide new insights into the molecular mechanisms of cardiac aging and suggest that targeting these pathways could offer therapeutic opportunities to address aging-related cardiac dysfunction.

The increase in IFN signatures across multiple cell types in the aging heart is particularly noteworthy. Our data indicate that interferon signaling plays an important role in modulating cell–cell communication within the aging heart. Using PopComm, we quantified ligand-receptor interactions and found that IFN-related pathways significantly influence these interactions, potentially altering the cellular microenvironment in aging hearts. This modulation of cell–cell communication could contribute to chronic inflammation and impaired tissue repair in the aging heart. Targeting these pathways could represent a novel approach to enhancing cardiac regeneration and reducing fibrosis in aging patients.

While our study presents important findings, there are several limitations that need to be addressed in future research. First, although our dataset is large, it is compiled from multiple studies, and batch effects significantly obscure biological signals, particularly subtle aging-related effects. The integration of diverse datasets introduces technical noise, which may hinder the identification of fine-grained biological patterns. Second, the age distribution of the samples we used is unbalanced, with relatively few young samples (<40 years old) included. To address these limitations, future studies should incorporate larger and more balanced cohorts across a broader range of ages, genders, and health statuses. Additionally, employing more advanced data integration techniques could help reduce batch effects and provide a clearer picture of the biological processes underpinning aging and disease.

This study provides critical insights into the cellular and molecular mechanisms underlying cardiac aging and disease. The identification of *NRG1*-expressing endocardial cells and the role of interferon signaling in aging suggest that these pathways could serve as potential targets for therapeutic intervention. Further investigation into these mechanisms may lead to the development of novel treatments for cardiac fibrosis and aging-related heart diseases. Additionally, PopComm is a useful framework for studying ligand-receptor interactions on the population scale, with potential applications beyond cardiac research. Future studies should explore the clinical utility of targeting interferon signaling and *NRG1*-expressing cells in the treatment of heart diseases, particularly in aging populations. Furthermore, the continued expansion of datasets with more diverse samples and the refinement of integrative analysis techniques will deepen our understanding of the heart’s cellular landscape and improve the precision of future cardiac research.

## Methods

### Data collection and metadata curation

We first retrieved the datasets from 12 studies covering both healthy and diseased hearts from our previously published DISCO database (https://www.immunesinglecell.org)^42,43^. Following stringent quality control (QC) procedures (see data preprocess section), 436 non-fetal cardiac samples from the 12 datasets were retained for integration (scRNA-seq, *n* = 79; snRNA-seq, *n* = 357). All datasets in this study were generated using 10x Genomics single-cell sequencing, reducing batch effects associated with different sequencing platforms^3,17–20,22,24,26,28,29,34,36^. Additionally, a spatial transcriptomics dataset acquired with the 10x Genomics Visium platform^19^ (**Supplementary Table 2**) was downloaded to validate the results of our PopComm method.

To obtain comprehensive clinical metadata for the samples, we manually curated information from the original data deposition databases, supplementary materials in published literature, and related documentation available on GitHub. The primary metadata variables included sample type (disease/normal), age, sex, disease diagnosis, sampling source (nuclei/cells), and tissue region. For some samples, additional variables such as body mass index (BMI), hypertension (HTN), diabetes, chronic kidney disease (CKD), smoking status, dyslipidemia, and the presence of heart failure were also incorporated. To ensure consistency and address recording discrepancies in certain project datasets, we standardized the key variables, including age group and tissue region. The original ages provided in the datasets were recorded and additionally categorized into 10-year bins to standardize the data, as some studies reported only age ranges. The reported heart regions were categorized into six standardized groups: left atrium (LA), right atrium (RA), left ventricle (LV), right ventricle (RV), apex (AP), and septum (S) to streamline downstream analysis and facilitate comparison across studies. Additional details about the sample metadata are provided in **Supplementary Table 2**.

### Data preprocess and quality control

From the DISCO database, we downloaded the gene expression matrices previously obtained by raw reads alignment to the human reference genome (hg38/GRCh38) and expression quantification with 10x Genomics Cell Ranger (v7.0.0)^76^ and the GENCODE annotation (v32). Subsequently, the gene expression matrix of each sample was further processed and analyzed using the R package “Seurat” (version 4.4.0)^77^. Low-quality cells were excluded based on the following criteria: mitochondrial read content >10%, nFeature_RNA <500, and nCount_RNA <500. For samples in the ERP123138 dataset, the mitochondrial read threshold was adjusted to 30% to account for dataset-specific characteristics. Subsequently, “scDblFinder” package (version 1.17.2, https://plger.github.io/scDblFinder/) was used to identify potential doublets/multiplets^78^. The detailed cell number information after each QC step is provided in **Supplementary Tables 3 and 4**. For the spatial transcriptomics data, the processed data provided by original publication were adopted.

### Data integration and cell type annotation

A hierarchical data integration strategy was employed to process the post-QC clean data, with single-cell and single-nucleus datasets integrated separately. Two rounds of integration were performed to identify broad and fine cell types. The integration tool employed was scVI^44^, a deep learning-based variational inference method. In the first round of integration, 5,000 highly variable genes across samples were selected as input to generate 30 batch-removed latent dimensions, enabling robust downstream analysis. The “FindNeighbors” function from the Seurat package was then used to construct a shared nearest neighbor graph, followed by the “FindClusters” function for unsupervised clustering. Cluster markers were identified using a one-versus-rest Wilcoxon rank-sum test via the “wilcoxauc” function provided by the presto package^79^ on Seurat-normalized expression; genes with a Benjamini-Hochberg (BH)-adjusted *P*-value < 0.01 and |log2FC| > 0.5 were retained, with area under the curve (AUC) and the detection-rate difference (|pct_in − pct_out|) used for ranking. For snRNA-seq, we downsampled to 50,000 nuclei to reduce computational burden and mitigate extreme cell-number imbalance across cell types. The cluster markers then were cross-checked against DISCO for annotation, and clusters belonging to the same broad cell types were merged.

We then conducted the second round of integration and annotation within each broad cell type. Here, we used 2,000 feature genes as input to scVI to generate 10 batch-corrected latent dimensions, as the heterogeneity within broad cell types is typically lower. The reduced number of features and dimensions also facilitated more precise analysis. For fine cell type annotation, additional manual processing was applied. Identified markers were carefully examined and compared with findings from previous studies to ensure accurate identification and classification of cell types.

### Phenotype association analysis

Phenotype association analysis was performed using snRNA-seq samples. For the analysis of aging, sex, and heart region, only the normal samples were included. For disease-related comparisons, the non-atrial samples were used to control for regional effects. The calculation of three distinct types of cell type proportions was conducted for each sample: broad cell type proportions, fine cell type proportions, and the percentage of each fine cell type within its corresponding broad cell type.

One-way Analysis of Variance (ANOVA) was employed to assess the disparities in cell type proportions across various phenotype groups. For the aging analysis, comparisons were made between the old (>60 years) and young (<60 years) groups. For sex, female and male samples were compared. The disease category comprised five comparisons: all disease samples versus normal samples, HF* versus normal, dilated cardiomyopathy (DCM) versus normal, arrhythmogenic cardiomyopathy (ACM) versus normal, and cardiac allograft vasculopathy (CAV) versus normal. HF* includes heart failure samples as well as DCM and ACM samples diagnosed with heart failure. For the heart regions, the comparisons were Ventricle versus Atrium, Left versus Right, and AP versus Ventricle. In each specific comparison, any related covariates were included in the model to control for potential confounding effects. The analysis was conducted using the “aov” function in R.

To determine whether a specific cell type was enriched or depleted in a given phenotype, the odds ratio (OR) was calculated based on a contingency table, where the rows represent “cells of the specific type” and “cells of other types,” while the columns represent “samples with the phenotype” and “samples without the phenotype.” The OR provided a quantitative measure of the relative abundance of the cell type within the phenotype compared to its abundance outside the phenotype.

For visualization, the OR values were scaled to the range of –5 to +5, where values <1 were transformed by –(1/OR). Adjusted *P*-values were calculated using the BH method and capped at 1×10^-15^. Only associations with an adjusted *P*-value < 0.05 were considered significant and included in the plot. The –log_10_ adjusted *P*-value and the |scaled OR| were used to define the color and size of the dots, respectively, while filled and hollow circles indicate enrichment and depletion patterns, respectively.

### Differentially expressed genes (DEGs) analysis

Two categories of DEGs were identified in the present study. The first category is cell type DEGs which were identified using the “wilcoxauc” function provided by the presto package^79^. Genes with a BH-adjusted *P*-value < 0.01 were retained for analysis.

The second category of DEGs analyzed was aging-related DEGs. We tested two different methods:

Method 1: The “FindAllMarkers” function from the Seurat package was used with the MAST method to identify DEGs between phenotype groups within specific cell types. MAST accounts for the fraction of genes expressed and allows the inclusion of covariates. In our analysis, sample identity was included as a covariate to mitigate the influence of nuclei correlation within samples. Additionally, metadata such as batch, sex, and heart region were incorporated as covariates to account for their inter-relationships with aging. DEGs were considered significant if they met the threshold of an adjusted *P*-value < 0.01.

Method 2: Pseudo-bulk data were generated for each cell type in each sample, and DESeq2 was employed to identify phenotype-specific DEGs^80^. Due to the limited sample size, covariates were not included in the analysis. DEGs were considered significant if they met the threshold of an adjusted *P*-value < 0.01.

We compared the results of these two methods and found a substantial number of overlapping DEGs (**Supplementary Fig. 4b; Supplementary Table 14 & 16**), demonstrating that both approaches consistently identified aging-related DEGs. For downstream analysis, the results from Method 1 were selected, as it incorporated covariates and thus provided a more robust framework.

### Analysis of gene expression dynamics along chronological age

For our analysis of the two largest datasets (EGAS00001006374^24^ and ERP123138^3^), the average normalized gene expression of each gene in each cell type was first calculated. Linear regression was then performed on the averaged expression levels, with chronological age as the independent variable and sex as a covariate within each dataset. Genes with a *P*-value < 0.01 were identified as aging-related genes. Gene module scores representing the expression of age-positive and age-negative gene sets were calculated for each cell using Seurat’s “AddModuleScore” function. Their association with age was then evaluated.

### PopComm

To quantify cell–cell communication at the population level, we developed PopComm, a computational framework that infers LR interactions across large-scale single-cell and single-nucleus RNA-seq datasets. PopComm enables sample-level communication scoring, making it suitable for correlating interaction strength with phenotypic variables such as age or disease status. PopComm consists of three main modules:

1. The LR Filtering Module identifies expressed ligand–receptor pairs between sender and receiver cell types, based on expression thresholds, correlation across samples, and linear modeling. The implementation of filtering can be applied to specific cell-type pairs or across all possible combinations.
2. The LR Scoring Module quantifies communication intensity at the sample-level using a projection-based scoring approach, allowing downstream comparison of communication strength across different phenotypes.
3. The Visualization Module provides functions to visualize LR interactions as circular networks, dot plots, heatmaps, PCA projections, and box/violin plots, supporting both cell type–level and sample-level interpretation of communication patterns.

The complete implementation and usage details are provided in the Application Note. Specific algorithmic details of PopComm are described in Supplementary Note “PopComm Methods” (Supplementary Materials), and the source code with documentation is available at https://github.com/JusticeGO/PopComm.

### Pathway analysis

Two types of pathway analysis were applied in this study. In scenarios where only a gene set was available, we used the “enrichGO” function from the clusterProfiler package^81^ to enrich Gene Ontology (GO) Biological Process terms, identifying the biological processes associated with a given gene set.

In cases where a ranked gene list was available, such as those ranked by correlation or other analyses, Gene Set Enrichment Analysis was performed using the fgsea package. This approach utilized the hallmark gene sets from MSigDB^82^ to uncover enriched pathways based on the ranking of genes.

In both cases, only pathways with an adjusted *P*-value of less than 0.05 were retained for further analysis.

### Gene set score

For certain analyses, we calculated scores for specific gene sets (e.g., the cell aging-related gene set) using the “AddModuleScore” function from the Seurat package. The gene set score generated by this function served as a surrogate for average expression, adjusted by subtracting the aggregated expression of control feature sets, which were set to 50 in this study. Specifically, for the IFN response gene set, we used the following genes: *IFI6*, *IFI44L*, *IFI44*, *IFI16*, *IFIH1*, *GRIFIN*, *IFITM5*, *IFITM2*, *IFITM1*, *IFITM3*, *IFITM10*, *IFIT2*, *IFIT3*, *IFIT1B*, *IFIT1*, *IFIT5*, *IFI27L1*, *IFI27*, *IFI27L2*, *IFI35*, *IFI30*, *ISG15*, *ISG20L2*, and *ISG20*.

### Cell type annotation of spatial data

The paired snRNA-seq data corresponding to the 10x Genomics Visium spatial transcriptomics data were downloaded from the original study. Broad cell types were annotated using the same approach as described above. Marker genes for each cell type were then identified. For the spatial data, cell type signatures at each spot were calculated using the identified marker genes and the “AddModuleScore” function. Within each slide, a spot was classified as a given cell type if its corresponding cell type signature exceeded the 90th percentile of a fitted normal distribution. For cardiomyocytes, due to their dominant proportion in the spatial data, a more relaxed threshold (above the 10th percentile) was applied. Each spot could be assigned to multiple cell types.

### Data visualization

Most figures were generated using visualization functions from the Seurat package. Additional plots were produced using ggplot2, ggpubr, pheatmap, and other commonly used R visualization packages.

## Supporting information

Supplementary Note for PopComm Methods

## Data availability

The raw data for each sample is available for download from the DISCO database (https://www.immunesinglecell.com). The integrated scRNA-seq and snRNA-seq datasets can be accessed via Zenodo at https://zenodo.org/records/17624791. The results of the PopComm analysis are also accessible via Zenodo at https://zenodo.org/records/16872202.

## Code availability

The key analysis code for this study is provided in (https://github.com/JusticeGO/HumanHeartAtlas_PopComm). The PopComm package is available on both PyPI (https://pypi.org/project/PopComm) and CRAN (https://CRAN.R-project.org/package=PopComm). PopComm’s further documentation and development resources are accessible via its GitHub repository (https://github.com/JusticeGO/PopComm).

## Acknowledgements

The research was supported by A*STAR under its BMRC Central Research Fund (CRF, UIBR) Award; AI, Analytics and Informatics (AI3) Horizontal Technology Programme Office (HTPO) seed grant (Spatial transcriptomics ST in conjunction with graph neural networks for cell–cell interaction #C211118015) from A*STAR, Singapore; Open Fund Individual Research Grant (Mapping hematopoietic lineages of healthy and high-risk acute myeloid leukemia patients with FLT3-ITD mutations using single-cell omics #OFIRG18nov-0103) from Ministry of Health, Singapore; National Research Foundation (NRF), Award no. NRF-CRP26-2021-0001; the National Research Foundation, Singapore, and Singapore Ministry of Health’s National Medical Research Council under its Open Fund-Large Collaborative Grant (“OF-LCG”) (MOH-OFLCG18May-0003); Singapore National Medical Research Council (#NMRC/OFLCG/003/2018); State Key Program of the National Natural Science Foundation of China (#82030059); National Natural Science Regional Innovation Fund joint fund key support projects (#U23A20485); Key R&D Program of Shandong Province (#2022ZLGX03); Taishan Pandeng Scholar Program of Shandong Province (#tspd20240819); Shandong Province Natural Science Foundation Youth Branch (#ZR2023QC209, #ZR2023QC313).

## Authors’ contributions

J.C. and Y.C. conceived and supervised the study. Z.G. and M.L. collected and curated the single-cell and single-nucleus datasets, performed the data analyses, and developed the PopComm framework. Z.G. and M.L. wrote the initial manuscript. K.S.A., Y.W., D.J.H., X.W., C.J.R., and X.Z. contributed to the biological interpretation of key findings and provided critical comments on the manuscript. K.S.A. further contributed essential revisions and refinement of the manuscript. All authors reviewed and approved the final manuscript.

