## Supplementary Information Notes for "A Population-Scale Single-Cell Atlas of the Human Heart Reveals Cellular Remodeling and Cell–Cell Communication in Aging and Cardiac Disease"

### Supplementary figure legends

**Supplementary Figure 1. Overview of fine cell type diversity and molecular signatures across scRNA-seq and snRNA-seq data.**

Integrated analysis of scRNA-seq and snRNA-seq datasets revealed the diversity and molecular signatures of fine-grained cell types. UMAP representations of the fine cell types in the integrated scRNA-seq and snRNA-seq data (left). Bar plots summarizing the relative proportions of fine cell types within each broad category (center). Dot plots highlight the top marker genes for each fine cell type, with color intensity indicating the expression level and dot size representing the percentage of expressing cells (right). For each fine cell type, the top three markers were selected based on fold change.

**Supplementary Figure 2. Association between myeloid subsets from scRNA-seq and snRNA-seq data.**

Heatmap depicting the strength of association between marker genes identified in the scRNA-seq and snRNA-seq data across myeloid cell subsets.

**Supplementary Figure 3. UMAP visualization of *NRG1*-expressing endocardial cells in scRNA-seq and snRNA-seq data, and in myocardial infarction and normal regions.**

- **a)** UMAP visualization of *NRG1*-expressing endocardial cells identified in the scRNA-seq and snRNA-seq data, grouped by disease status (HF* vs. Normal). HF* includes heart failure samples as well as DCM/ACM samples diagnosed with heart failure.
- **b)** UMAP visualization of NRG1-expressing endocardial cells in an independent dataset containing myocardial infarction (MI) samples. Feature plots, from left to right, of NRG1 expression, categorization by disease status (MI vs. normal), and anatomical region (FZ, IZ, BZ, RZ, and normal).

**Supplementary Figure 4. Supplementary analysis of aging-related gene expression changes in various cell types.**

- **a)** Bar plot showing the total number of aging-related DEGs (positive and negative) per fine cell type in the snRNA-seq data, Numbers indicate upregulated (red) and downregulated (blue) DEGs in old vs. young comparison.
- **b)** Venn diagram showing the overlap of aging-related DEGs between Seurat's MAST and Pseudo-bulk methods.
- **c)** Boxplots showing *NPPA* expression in different ages and heart regions.
- **d)** Boxplots showing the interferon-induced gene expression score across cell types, grouped by broad and fine categories. Statistical significance was assessed using the Kruskal-Wallis test.
- **e)** Aging-related DEGs identified in the scRNA-seq data across major cell types.

  Left panel: Bar plot showing the total number of aging-related DEGs (positive and negative) per cell type. Numbers indicate upregulated (red) and downregulated (blue) DEGs in old vs. young comparison.

  Right panel: Upset plot illustrating overlaps of aging-related DEGs among cell types. Bars show the number of shared DEGs across combinations of cell types, with the colors indicating directionality consistency: positively consistent (red), negatively consistent (blue), or inconsistent (gray).

**f)** Heatmap showing the expression patterns of aging-related DEGs shared across multiple cell types in the scRNA-seq data. Each row represents a gene, and each column represents a cell type. Color intensity indicates the log2 fold change (old vs. young), with red indicating upregulation and blue indicating downregulation.

**Supplementary Figure 5. Supplementary analysis of population-scale ligand-receptor interactions identified by PopComm across cell types.**

**a)** Correlation between the number of ligand-receptor (LR) pairs identified and the number of individuals in which each cell type was detected.

**b-c)** Dot plots showing the top 5 ligand–receptor interactions per sender–receiver pair in both scRNA-seq (b) and snRNA-seq (c) datasets. Dot size indicates the correlation coefficient across individuals and color intensity represents the adjusted *P*-value of the interaction.

### Supplementary tables

**Supplementary Table 1. Summary of published single-cell transcriptomic studies of the human heart.**

**Supplementary Table 2. Metadata of human heart samples used in this study.**

**Supplementary Table 3. Quality control metrics for individual samples.**

**Supplementary Table 4. Aggregated quality control metrics by project.**

**Supplementary Table 5. DEGs of broad cell types in scRNA-seq data.**

**Supplementary Table 6. DEGs of fine cell types in scRNA-seq data.**

**Supplementary Table 7. DEGs from fine vs. broad cell types comparisons in scRNA-seq data.**

**Supplementary Table 8. DEGs of broad cell types in snRNA-seq data.**

**Supplementary Table 9. DEGs of fine cell types in snRNA-seq data.**

**Supplementary Table 10. DEGs from fine vs. broad cell types comparisons in snRNA-seq data.**

**Table S5-S10 Notes:**

**Adjusted *P*-value < 0.01**

**feature:** The gene identifier.
**group:** The cell type or cell subtype where the gene is expressed.
**avgExpr:** The average expression level of the gene across the cells in the group.
**Log2FC:** Log2 fold change of gene expression between conditions.
**statistic:** The test statistic used to evaluate the significance of the differential expression.

| | |
|---|---|
| **auc:** | The area under the ROC curve, representing the ability of the gene to distinguish between the groups. |
| **pval:** | The *P*-value from the statistical test. |
| **padj:** | The adjusted *P*-value after controlling for multiple tests. |
| **pct_in:** | The percentage of cells expressing the gene in the cell type of interest. |
| **pct_out:** | The percentage of cells expressing the gene in other cell types. |
| **pct_diff:** | The difference in gene expression percentage between the cell type of interest and other cell types. |
| **compare_to:** | The reference broad cell type for the comparison (for fine vs. broad comparisons). |

**Supplementary Table 11. Cell type-phenotype associations across broad and fine resolutions.**

**Supplementary Table 12. Spearman correlation between sample-level cardiomyocyte gene expression and the proportion of NRG1-expressing endocardial cells (*P*-value < 0.05).**

**Supplementary Table 13. Aging-related DEGs identified from scRNA-seq data at broad cell type level (adjusted *P*-value < 0.01).**

**Supplementary Table 14. Aging-related DEGs identified from snRNA-seq data at broad cell type level (adjusted *P*-value < 0.01).**

**Supplementary Table 15. Aging-related DEGs identified from snRNA-seq data at fine cell type level (adjusted *P*-value < 0.01).**

**Supplementary Table 16. Aging-related DEGs identified from snRNA-seq pseudo-bulk data at broad cell type level with DESeq2 (adjusted *P*-value < 0.01).**

**Supplementary Table 17. Significant ligand-receptor-sender-receiver pair results for scRNA-seq data by PopComm (adjusted *P*-value < 0.05).**

**Supplementary Table 18. Significant ligand-receptor-sender-receiver pair results for snRNA-seq data by PopComm (adjusted *P*-value < 0.05).**

**Supplementary Table 19. Significant ligand-receptor-sender-receiver pair results for control snRNA-seq data by PopComm (adjusted *P*-value < 0.05).**

**Table S17-S19 Notes:**

**Adjusted *P*-value < 0.05**

| | |
|---|---|
| **ligand, receptor:** | Ligand and receptor gene symbols. |
| **cor:** | Correlation coefficient. |
| **p_val:** | Raw *P*-value. |
| **adjust.p:** | Adjusted *P*-value. |
| **sender, receiver:** | Sender and receiver cell types. |
| **slope:** | Slope of the linear regression model. |
| **intercept:** | Intercept of the linear regression model. |

**Supplementary Table 20. Significant differentially active ligand–receptor (DALR) pair results for snRNA-seq data when comparing between disease and control (adjusted *P*-value < 0.01).**

**Supplementary Table 21. Significant differentially active ligand–receptor (DALR) pair results for control snRNA-seq data when comparing between IFN type high and low (adjusted *P*-value < 0.01).**

**Supplementary Table 22. Significant differentially active ligand–receptor (DALR) pair results for control snRNA-seq data when comparing between age group old and young (adjusted *P*-value < 0.01).**

**Table S20-S22 Notes:**

| | |
|---|---|
| **Adjusted *P*-value < 0.01** | |
| **DALR-SR:** | Differentially active ligand-receptor-sender-receiver pair |
| **Estimate:** | Estimate of the regression model. |
| **Std_error:** | Standard error of the estimate. |
| **t_val:** | t-statistic. Tests whether the estimate is significantly different from zero. |
| **p_val:** | Raw *P*-value. |
| **adjust.p:** | Adjusted *P*-value. |
| **ligand, receptor:** | Ligand and receptor gene symbols. |
| **sender, receiver:** | Sender and receiver cell types. |
| **slope:** | Slope of the linear regression model. |
| **intercept:** | Intercept of the linear regression model. |
