## Supplementary figures and images for "A Population-Scale Single-Cell Atlas of the Human Heart Reveals Cellular Remodeling and Cell–Cell Communication in Aging and Cardiac Disease"

### Supplementary Figure 1

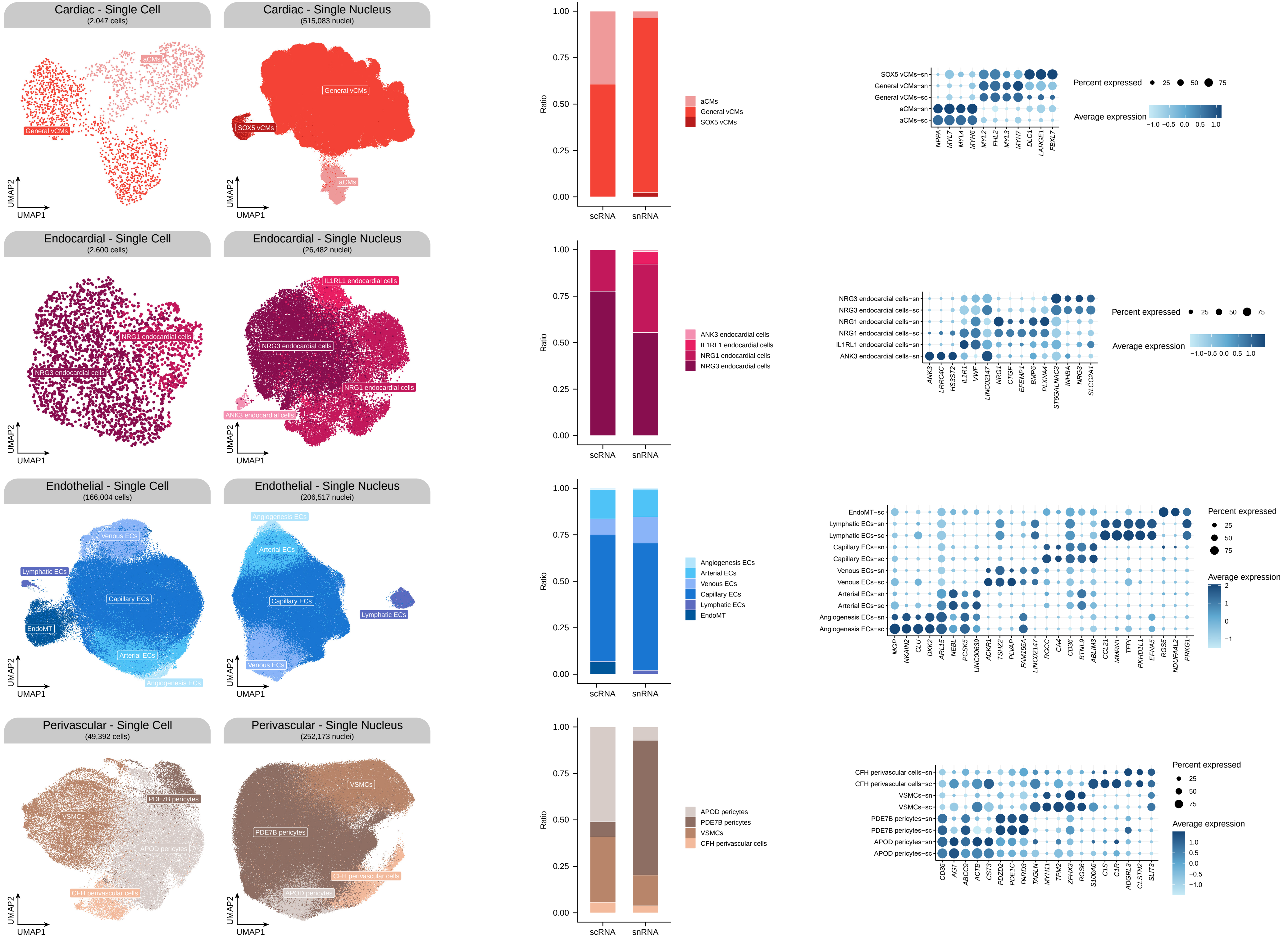

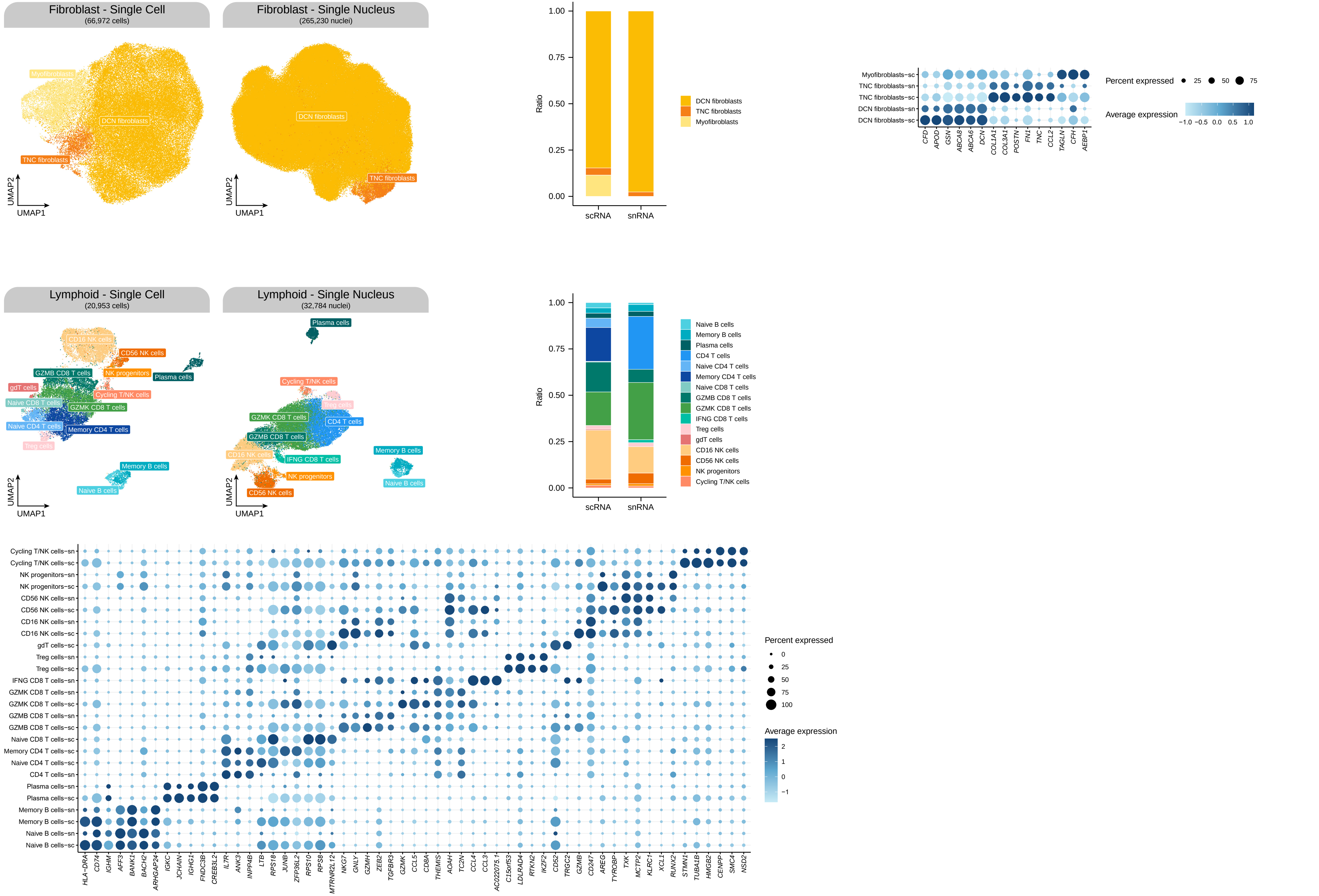

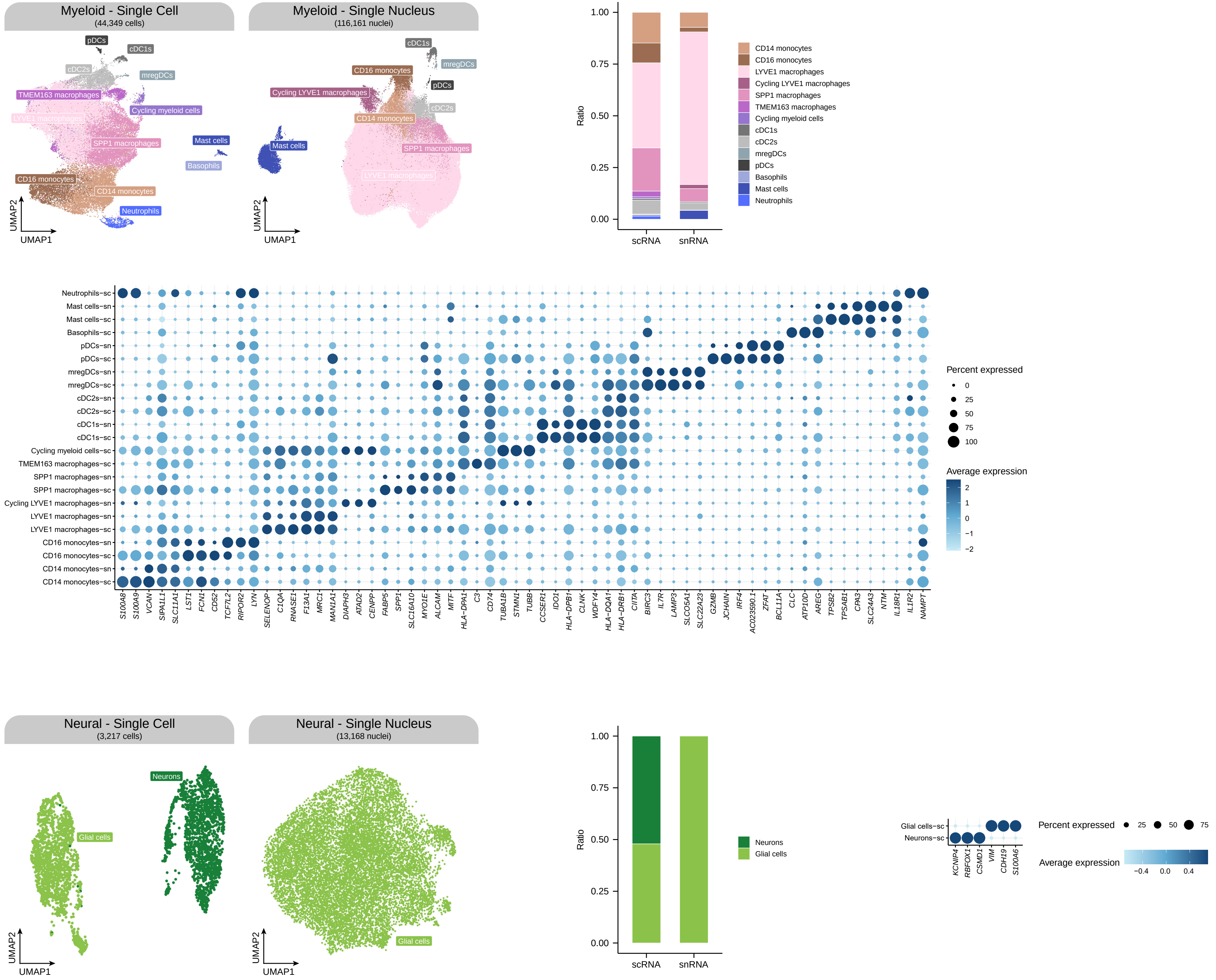

### Supplementary Figure 2

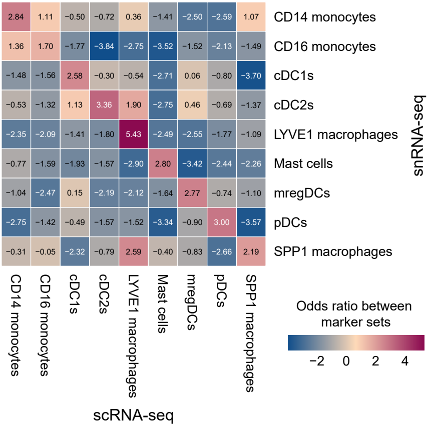

### Supplementary Figure 3

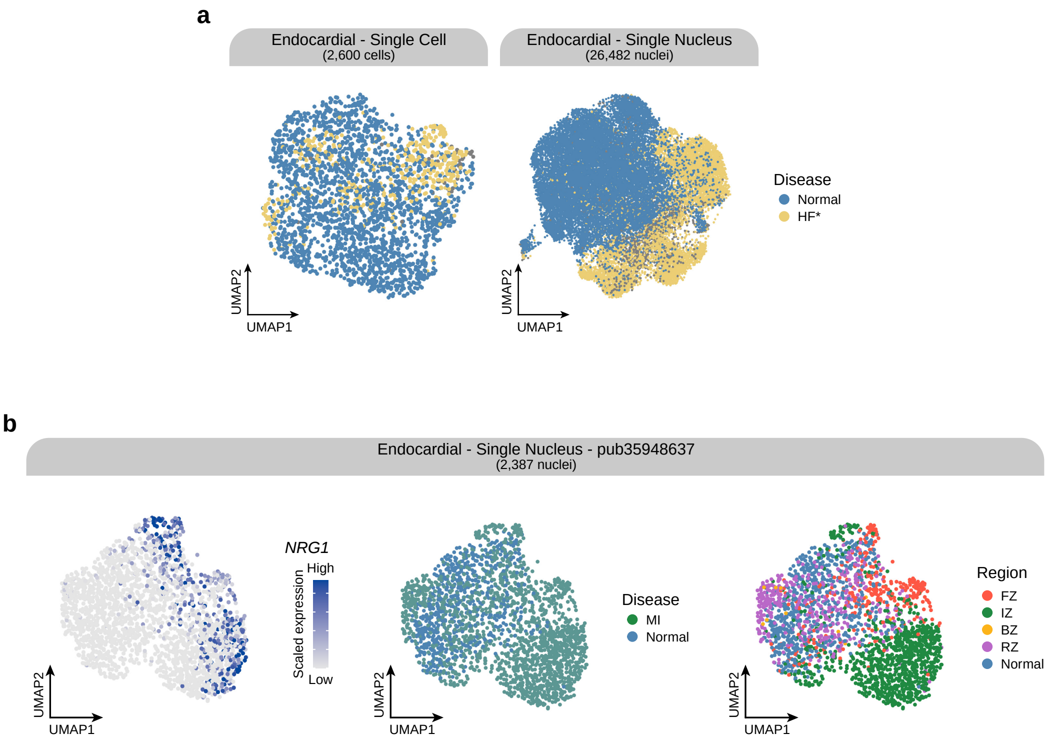

### Supplementary Figure 4

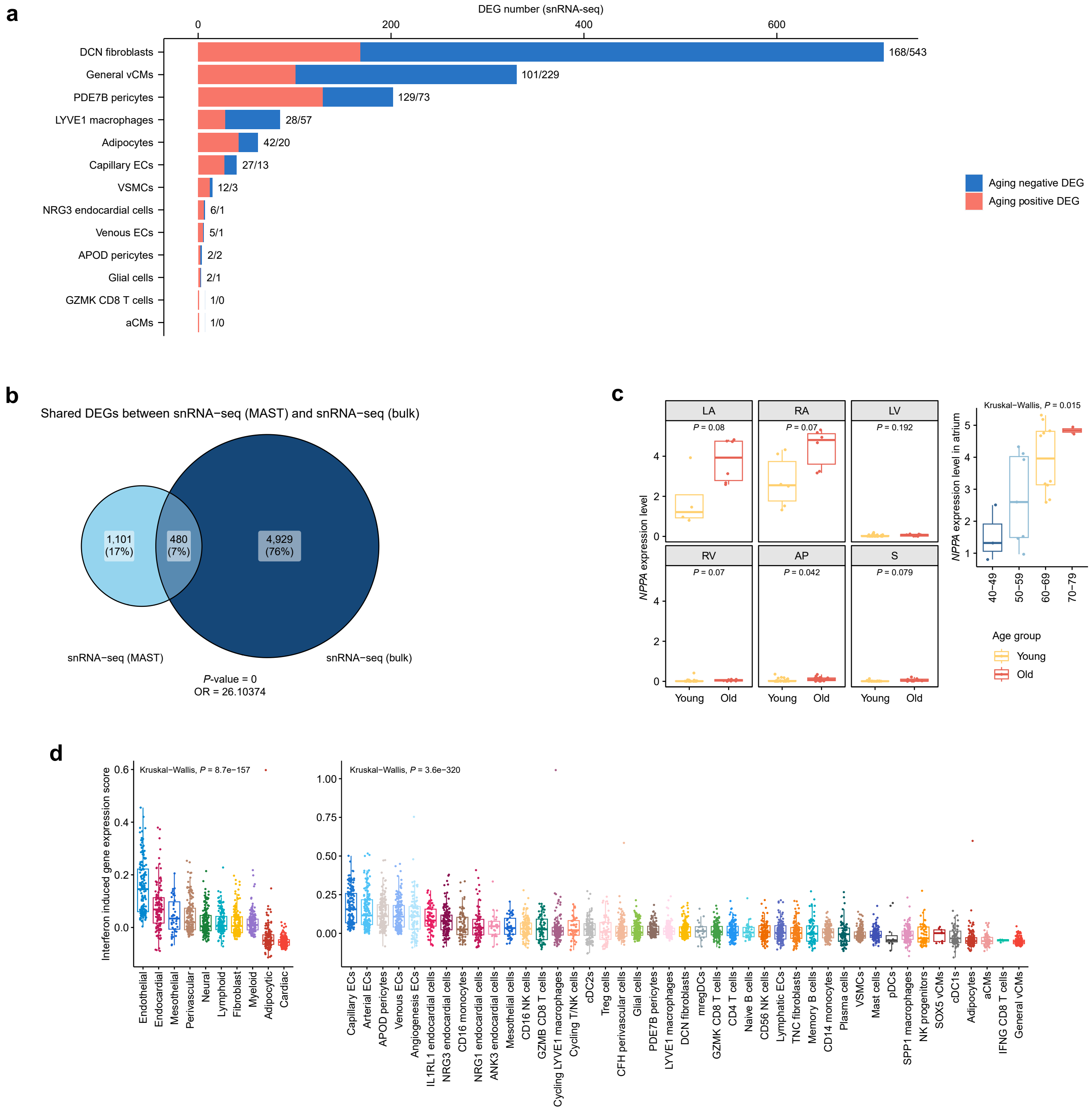

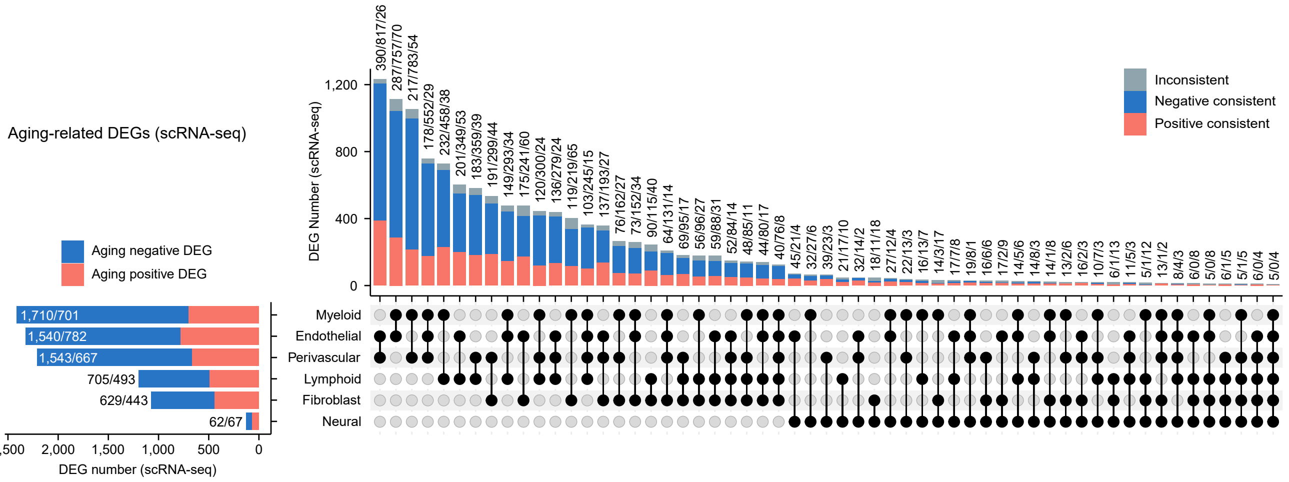

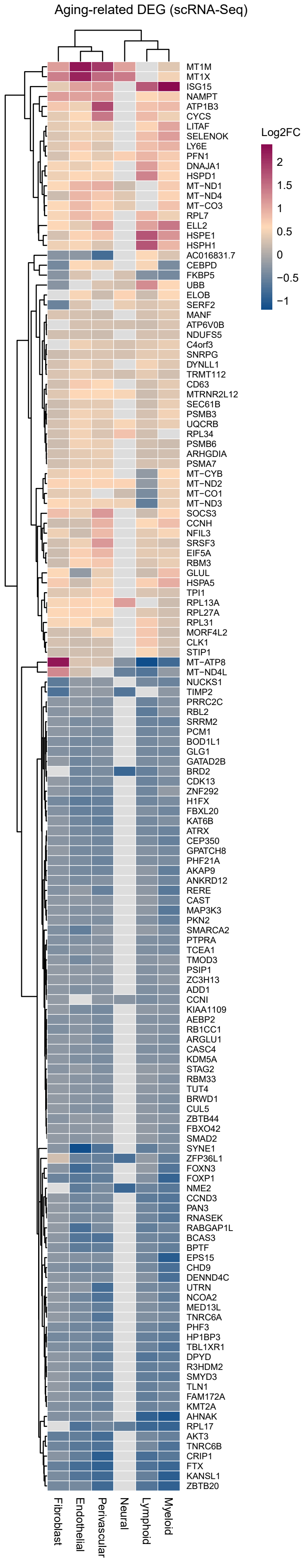

### Supplementary Figure 5

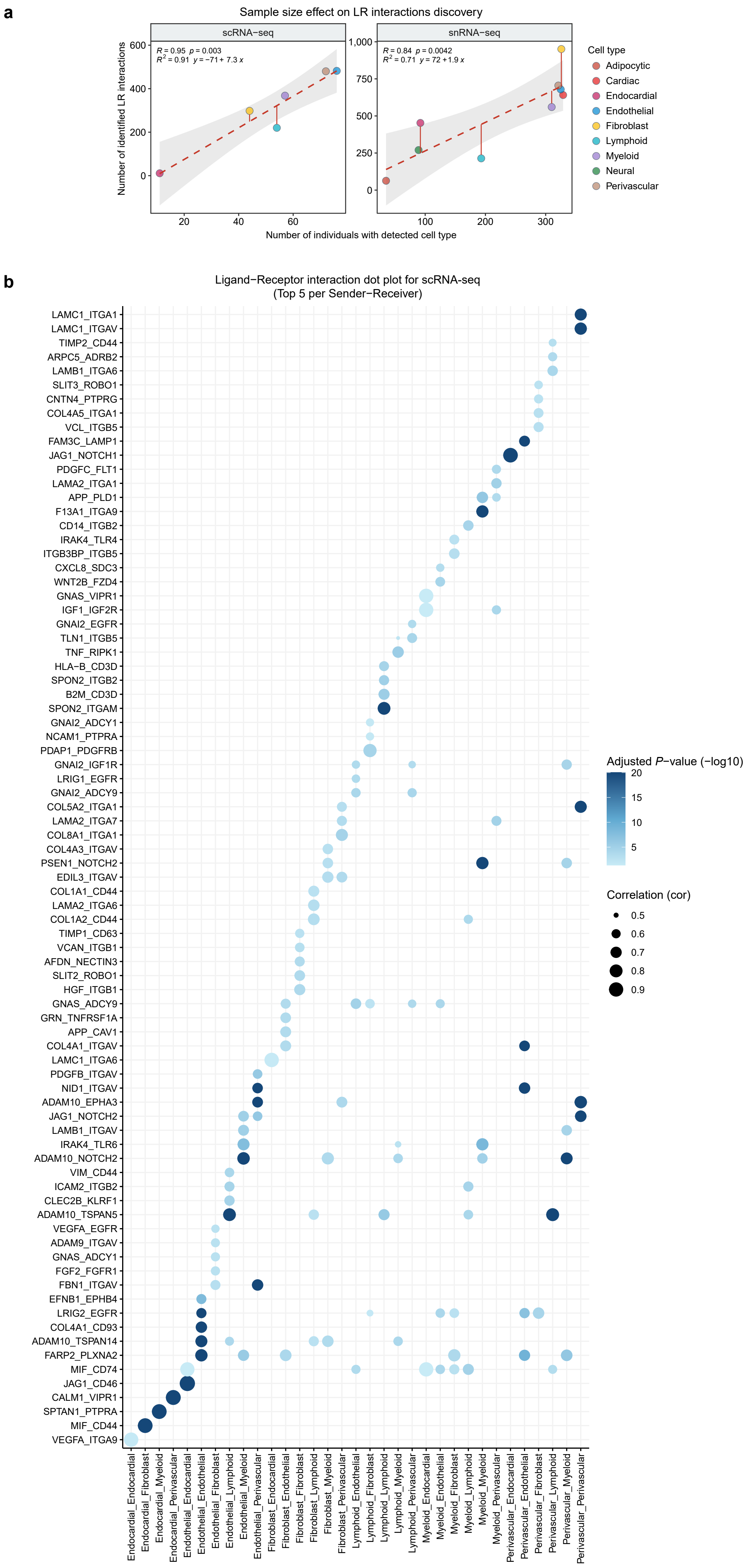

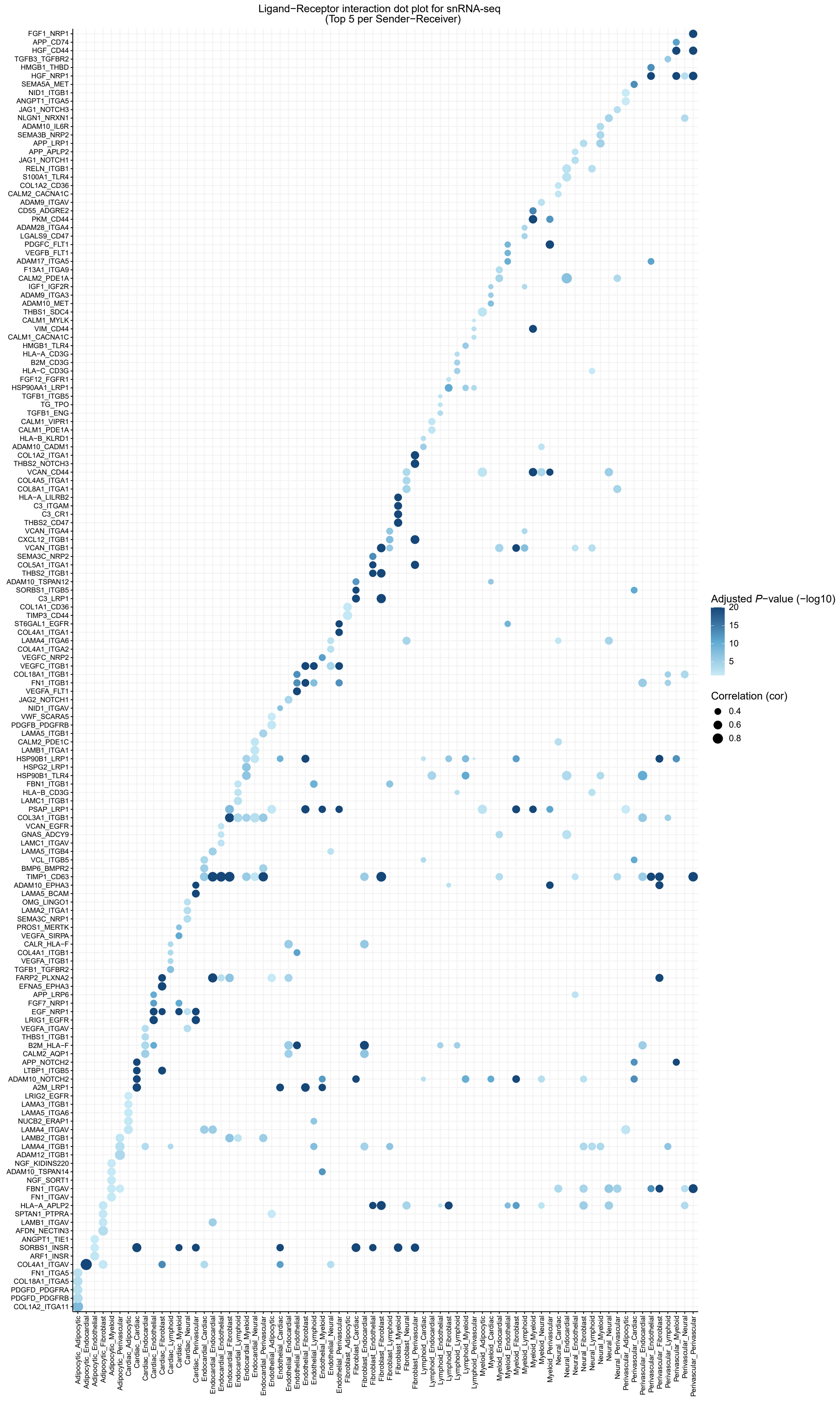
