## Supplementary Note for PopComm Methods for "A Population-Scale Single-Cell Atlas of the Human Heart Reveals Cellular Remodeling and Cell–Cell Communication in Aging and Cardiac Disease"

**Ligand-receptor (LR) database**

The LR database used in PopComm was primarily obtained from CellTalkDB (https://xomics.com.cn/celltalkdb/), a curated resource constructed through a combination of text mining and manual verification of protein–protein interaction databases. It contains 3,398 human LR pairs, each supported by published experimental evidence. In addition to CellTalkDB, PopComm also supports LR annotations from other databases, including CellChat and CellPhoneDB, offering flexibility for users with different reference preferences.

**LR Filtering Module**

To detect potential cell–cell communication events, PopComm includes a dedicated LR Filtering Module featuring two core functions: filter_lr_single() and filter_lr_all(). These functions filter ligand–receptor (LR) interactions by evaluating the expression-based correlation between ligands expressed in the sender cell type and receptors expressed in the receiver cell type across samples. This ensures that only biologically plausible and sample-supported LR interactions are retained for downstream analysis.

**Filtering LR for Specific Cell Type Pairs**

The filter_lr_single() function is designed to identify ligand–receptor (LR) interactions between a user-defined sender and receiver cell type by evaluating their expression patterns across multiple samples. The filtering process involves several key steps to ensure that only biologically meaningful and statistically supported interactions are retained.

First, the function performs sample filtering by selecting only those samples in which both the sender and receiver cell types contain more than a minimum number of cells, defined by min_cells (default: 50). To ensure robustness, it retains only cell type pairs that satisfy this condition in more than min_samples (default: 10) samples.

Next, a preliminary LR filtering step is applied. From the reference ligand–receptor database, the function filters LR pairs whose ligand and receptor genes are expressed in more than min_cell_ratio (default: 0.1) of cells within the sender and receiver populations, respectively. For these filtered genes, the average expression per sample–cell type pair is calculated.

The core of the analysis lies in expression correlation analysis. For each LR pair, the function calculates the correlation between ligand and receptor expression across samples using the method defined by cor_method (default: "spearman"). Statistical significance is assessed using cor.test(), and a linear model (lm(receptor ~ ligand)) is fitted to extract slope and intercept values. To mitigate the influence of extreme values, outlier removal is performed using the remove_outlier() function before the correlation analysis.

To be retained, LR pairs must meet all of the following criteria: a correlation coefficient greater than min_cor (default: 0), an adjusted *P*-value below min_adjust_p (default: 0.05), and co-expression of both ligand and receptor in more than min_sample_ratio (default: 0.1) of the retained samples. Finally, all *P*-values are corrected for multiple hypothesis testing using the method specified in adjust_method (default: "BH", i.e., Benjamini–Hochberg correction).

This rigorous filtering process ensures that only robust and biologically relevant LR interactions are passed on for scoring and downstream analyses.

**Filtering LR Across All Cell Type Combinations**

The filter_lr_all() function automates the process of identifying meaningful ligand–receptor (LR) interactions by systematically evaluating all possible sender–receiver cell type combinations, including self-interactions. For each pair, it internally calls the filter_lr_single() function to assess the expression-based correlation between ligand and receptor genes across samples.

To efficiently handle large datasets, the function supports multi-threaded execution on both Windows and Linux systems, with the number of parallel processes controlled via the num_cores parameter.

The final output is a structured DataFrame that compiles all retained LR pairs along with their associated sender and receiver cell types, correlation coefficients, statistical significance, and other relevant metadata. This filtered dataset serves as the foundation for downstream scoring, condition-based comparisons, and visualization of intercellular communication patterns.

**Ligand-Receptor Scoring Module**

To quantify the communication intensity of candidate LR pairs across samples, the Ligand–Receptor Scoring Module includes two functions—score_lr_single() and score_lr_all()—which compute sample-level interaction scores based on linear model projections in expression space.

**Scoring for Specific Cell Type Pair**

The score_lr_single() function quantifies the communication strength between a specified sender and receiver cell type across multiple samples. To ensure statistical reliability, the function first filters for samples in which both cell types have more than min_cells (default: 50) cells, consistent with the criteria used in the filter_lr_single() function.

For each retained sample, the function begins by calculating the average gene expression of each ligand and receptor within their respective cell types. These expression values are then matched to ligand–receptor pairs provided in the filtered_lr input, which typically comes from prior filtering steps such as filter_lr_single() or filter_lr_all().

Next, for each LR pair, the sample's ligand and receptor expression levels are projected onto a pre-fitted regression line defined by that pair's slope and intercept, capturing the expected relationship between the two genes. The raw interaction score is then computed as the Euclidean distance between the projected point and a baseline, which is defined by the sample with the minimum overall expression for that pair.

To enable comparison across samples and conditions, the raw scores are subsequently normalized to a 0–1 range using min–max scaling, resulting in a final normalized_score that reflects the relative communication intensity for each sample.

**Scoring Across All Cell Type Combinations**

The score_lr_all() function quantifies ligand–receptor communication strength across all sender–receiver cell type combinations. It begins by grouping the input filtered_lr table by sender, receiver, and LR pair, and then applies the score_lr_single() function to each group to compute sample-level interaction scores. This process ensures that every relevant interaction is evaluated in the context of all observed cell type pairs.

To improve computational efficiency, the function supports parallel execution, with the number of threads controlled by the num_cores parameter. This allows the scoring process to scale effectively across large datasets and multi-core systems.

The output is a comprehensive data frame that integrates multiple layers of information, including metadata from the original filtered_lr input (such as ligand, receptor, and cell type identity), the sample ID, and both the raw interaction score and its min–max normalized counterpart (normalized_score). Within each LR pair, samples are automatically ranked by communication intensity, which facilitates a range of downstream analyses, including group comparisons, phenotype associations, and visual exploration of inter-sample variability in communication patterns.

**Visualization Module**

To intuitively illustrate LR interactions and sample-level communication dynamics, PopComm offers several visualization submodules, grouped into two categories: LR Pair Visualization and Sample Score Visualization.

**LR Pair Visualization**

To visualize interaction patterns between sender and receiver cell types, PopComm offers two intuitive plotting functions: circle_plot() and dot_plot().

The circle_plot() function produces a circular network diagram that provides a global view of cell–cell communication. In this layout, nodes represent distinct cell types, and directed edges indicate the direction of communication from sender to receiver based on ligand–receptor interactions. The edge width is scaled according to interaction strength, such as the correlation coefficient or number of significant interactions, while the edge color encodes the overall interaction intensity. The function also supports optional display of self-interactions (e.g., autocrine loops), enhancing interpretability of intra-cellular signaling. Users can customize node colors or use predefined palettes. The output is returned as a ggplot object, making it fully modifiable for publication-quality figures.

The dot_plot() function creates a dot matrix plot to summarize LR interactions across multiple sender–receiver pairs. In this visualization, dot size corresponds to the strength of the interaction (e.g., correlation), and dot color reflects statistical significance, represented as the negative log10 of the adjusted *P*-value (capped at 1e–20 for stability). Users can highlight the top LR pairs per cell type combination using the top_n parameter or display specific interactions via the selected_LR option. The function also supports different scaling strategies for dot size, including absolute correlation or radius-based scaling. Like circle_plot(), this function returns a ggplot object for flexible customization.

Together, these two tools provide powerful and visually intuitive ways to explore the directionality and strength of intercellular communication across diverse cell populations.

**Sample Score Visualization**

PopComm provides several visualization tools to analyze sample-level variability in ligand–receptor (LR) communication, which is crucial for uncovering biological or clinical patterns across different conditions. These functions allow both unsupervised exploration and group-based comparisons using interaction scores derived from the score_lr module.

The heatmap_sample() function generates a clustered heatmap where rows represent ligand–receptor–sender–receiver (LRSR) pairs and columns represent individual samples. It accepts both interaction score data and corresponding sample metadata, and allows subsetting by specific sender or receiver cell types. Hierarchical clustering is applied to both rows and columns to reveal structure in the communication landscape, and column annotations can be added based on sample attributes such as treatment, age, or sex.

To further explore major axes of communication variation, the pca_sample() function performs principal component analysis (PCA) on sample-level LR scores. The results are projected into a two-dimensional PCA space to visualize overall sample clustering. Users can subset by specific sender or receiver types, and sample points can be colored by metadata attributes to highlight clustering driven by biological conditions or clinical variables.

For assessing group-level differences in communication strength, boxplot_lr_group_comparison() provides a straightforward visualization of a specific LRSR interaction across categorical sample groups. It displays a boxplot with jittered data points overlaid, enabling clear comparison of communication intensity between groups such as “young” and “aged.”

Finally, the dotplot_lr_continuous_group() function evaluates how LR interaction strength relates to a continuous phenotype such as age or disease severity. It creates a scatterplot of interaction scores versus the continuous variable, and optionally adds a linear regression line to assess correlation trends.

Together, these tools offer a comprehensive framework for visualizing and interpreting how cell–cell communication varies at the sample level across diverse biological contexts.

**Differential Expressed Interaction Module**

The Differential Expressed Interaction Module in PopComm is designed to uncover ligand–receptor (LR) interactions whose communication strength significantly varies across different biological or clinical conditions. In this analysis framework, the sample-level interaction score—calculated using the scoring module—is treated as the dependent variable, while the phenotype of interest (e.g., disease status, age group, or treatment) serves as the independent variable. Additional covariates such as age, sex, or batch can be incorporated to account for potential confounding effects.

For each ligand–receptor–sender–receiver (LRSR) pair, a linear regression model is fitted to examine the relationship between communication strength and the phenotype, with the option to adjust for covariates. The model estimates the effect of the phenotype on the LR score, and significance is assessed through *P*-values derived from the model fit. To correct for multiple testing across many LR pairs, Benjamini–Hochberg (BH) adjustment is applied to control the false discovery rate (FDR).

Only LR interactions with an adjusted *P*-value (FDR) below 0.05 are retained, representing interactions that are significantly different in strength between groups or correlated with a continuous phenotype. This module enables systematic discovery of condition-associated cell–cell communication changes, offering critical insights into how interaction patterns shift across developmental stages, disease progression, or treatment response.

**Implementation**

We have developed PopComm in both R and Python to ensure broad accessibility and integration with diverse analysis pipelines. The R version has been submitted to CRAN (https://CRAN.R-project.org/package=PopComm) and is available for installation using standard package management tools. Similarly, the Python version has been published to PyPI (https://pypi.org/project/PopComm/), allowing easy installation via pip. PopComm's further documentation and development resources are accessible via its GitHub repository (https://github.com/JusticeGO/PopComm).

Both implementations offer consistent functionality and share the same core logic, enabling users to seamlessly switch between languages based on their preference or project requirements. We are committed to long-term maintenance and regular updates of both versions, including feature enhancements, performance improvements, and compatibility with emerging single-cell and spatial transcriptomics platforms. Extensive documentation and usage examples will be provided to support users in deploying PopComm across a variety of biological contexts.
